# Structural and dynamic basis of indirect apoptosis inhibition by Bcl-xL: a case study with Bid

**DOI:** 10.1101/2025.08.31.673340

**Authors:** Christina Elsner, Anton Hanke, Oscar Vadas, Francesco L. Gervasio, Enrica Bordignon

**Affiliations:** Department of Physical Chemistry, University of Geneva, 1211 Geneva, Switzerland; Department of Pharmaceutical Sciences, University of Geneva, 1211 Geneva, Switzerland; Swiss institute of Bioinformatics, University of Geneva, 1211 Geneva, Switzerland; Department of Microbiology and Molecular Medicine, University of Geneva, 1211 Geneva, Switzerland; Chemistry Department, University College London, WC1E 6BT London, United Kingdom

**Keywords:** Apoptosis, Bid, Bcl-xL, DEER, molecular dynamics

## Abstract

Intrinsic apoptosis is a form of cell death which is activated, executed and inhibited by the Bcl-2 protein family. The structural basis of the inhibition mechanisms remains elusive. Here, we characterize the ensemble structural model of the inhibitory Bcl-xL/tBid complex at the mitochondrial membrane by probing inter-residue distances and dynamic solvent accessibilities complemented by integrative modelling and molecular dynamics simulations. We show that Bcl-xL and tBid form a heterodimer anchored to the membrane by the C-terminal helix of Bcl-xL. The BH3 domain of tBid is wedged between the exposed hydrophobic groove of Bcl-xL and the membrane headgroups, while tBid’s C-terminal helices remain dynamically engaged with the bilayer. This dynamic architecture sheds light on the mechanism of indirect inhibition of apoptosis.

**Significance Statement:** Programmed cell death, or apoptosis, is a fundamental process that eliminates damaged cells. However, cancer cells often evade this process by overexpressing anti-apoptotic Bcl-2 family proteins, which neutralise their pro-apoptotic counterparts. The structural details of the molecular mechanisms governing this inhibition have remained elusive and up to now, only partial structures could be obtained with high-resolution methods. In this study, we provide the structural model of the inhibitory complex formed by the anti-apoptotic protein Bcl-xL and the pro-apoptotic protein Bid at the mitochondrial membrane using experimental constraints and molecular dynamics simulations. Our work provides the structural basis of one of the inhibitory mechanisms of cell death, offering a new framework for developing strategies to overcome therapeutic resistance in cancer.

## Introduction

Apoptosis, a form of programmed cell death in multicellular eukaryotes, is a fundamental biological process; its dysregulation is implicated in the pathogenesis of numerous severe diseases [1–3]. The key step of the intrinsic pathway of apoptosis is the irreversible mitochondrial outer membrane permeabilization (MOMP), which is tightly controlled by the B-cell lymphoma 2 (Bcl-2)-protein family. Bcl-2 proteins are characterized by up to four Bcl-2 homology (BH) domains and, in most cases, a C-terminal transmembrane domain (TMD). Only the BH3 domain is common to all family members and essential for mediating their diverse interactions [4–7] (see also Fig. 1).

**Figure 1:**
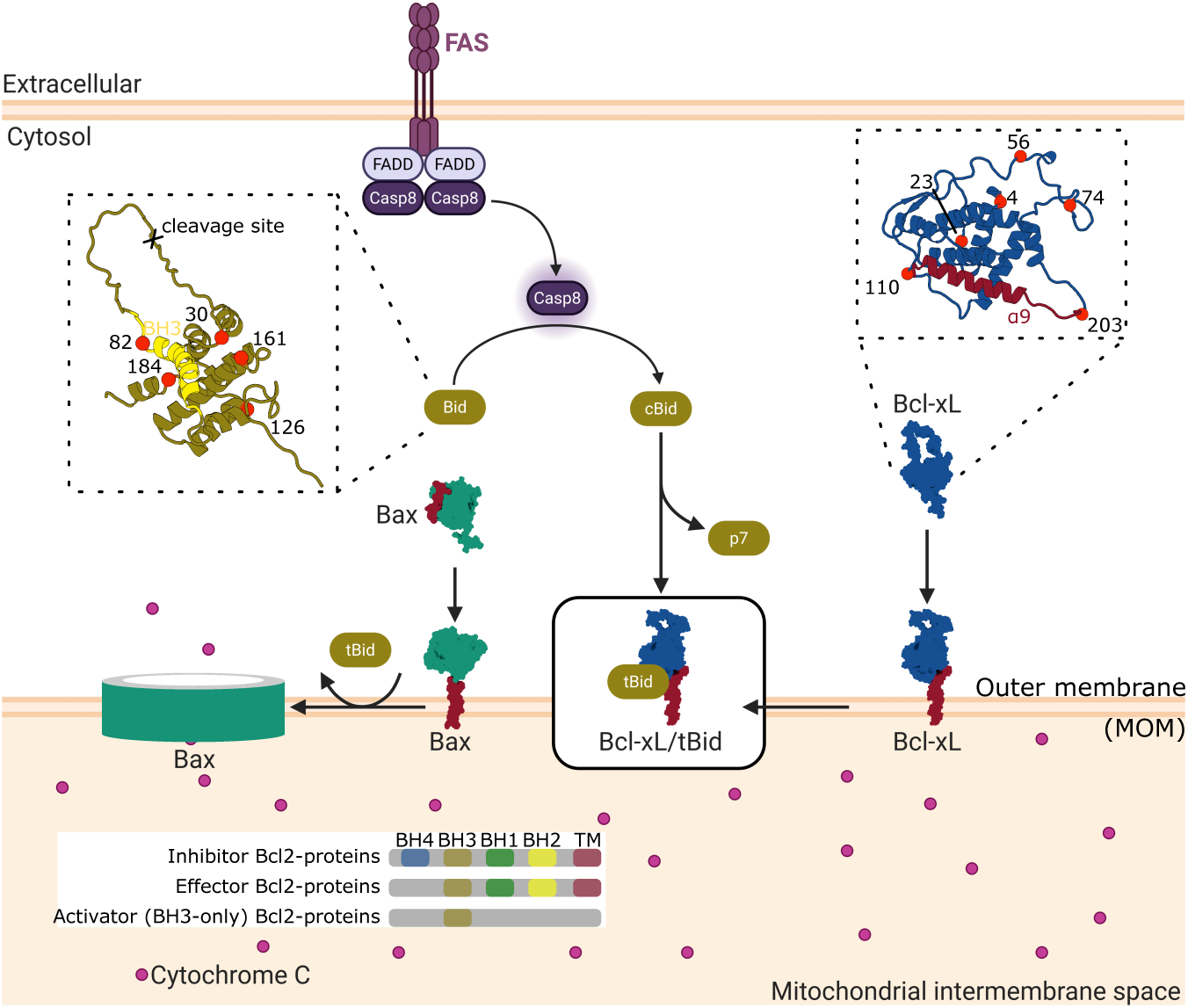
Scheme of the indirect inhibition of apoptosis in a minimal interaction network. External apoptotic stimuli activate Caspase 8, which cleaves Bid, producing cBid (consisting of the 7 kDa p7 fragment and the C-terminal 15 kDa tBid fragment). Monomeric Bax shuttles from the cytosol to the membrane via its C-terminal helix (dark red). At the membrane, the tBid fragment activates Bax, which forms oligomers permeabilizing the MOM (left). If tBid is sequestered by Bcl-xL at the membrane, membrane permeabilization is indirectly inhibited (right). The inset on the left shows the solution structure of mouse Bid (1DDB) with the helix containing the BH3 domain (yellow, BH3 domain: residues 87-100), the cleavage site in the loop and the spin labeling sites (red) indicated. The inset on the right shows the solution structure of full-length human Bcl-xL [21] with the C-terminal hydrophobic helix (dark red) and the spin labeling sites (red) indicated.

The Bcl-2 family is broadly divided into three subclasses. The first two are multi-BH-domain proteins: pro-apoptotic effectors (e.g., Bax, Bak, Bok) that oligomerize to permeabilize the mitochondrial outer membrane (MOM), and anti-apoptotic proteins (e.g., Bcl-2, Bcl-xL) that inhibit MOMP. The third subclass consists of the BH3-only proteins (e.g., Bim, Bid, Bad, PUMA), which act as either activators by directly triggering effectors or as sensitizers by neutralizing the anti-apoptotic proteins. These proteins dynamically exchange interaction partners, shuttling between the cytosol and the MOM in a process termed the “dance of death” [6]. This complex process involves substantial conformational changes and critically depends on interactions with the MOM (see for example [8–10]).

To date, structural studies have primarily focused on the oligomeric state of effector proteins at the membrane. While the transformation of Bax and Bak from soluble monomers to membrane-bound oligomers is partially understood (see for example [11–16]), and recent cryo-EM structures have provided new clues about pore assembly based on tetrameric units [17], the topology of these oligomers within the membrane remains unclear. More critically, despite their relevance as therapeutic targets [18], the atomistic structures of inhibitory complexes between full-length anti-apoptotic proteins and their effector or activator partners remain unresolved. While analyses of truncated, water-soluble proteins have revealed a canonical BH3-in-groove binding mode [10, 19–30], emerging evidence suggests that inhibitory interactions can involve more than one site [5, 31, 32]. This highlights the necessity of studying full-length proteins to fully elucidate the complex mechanism of inhibition.

Here, we investigate the molecular basis for Bcl-xL’s cytoprotective activity within a minimal, representative interaction network comprising the effector Bax, the activator cBid, and Bcl-xL itself (see Fig. 1). Using an integrative approach that combines modeling with extensive molecular dynamics (MD) simulations, constrained by available structural and topological data, we generated an atomistic model of the inhibitory Bcl-xL/tBid complex at the MOM which is validated by *in vitro* and *in organelle* data. The resulting structural ensemble reveals an inhibitory dimer with a distinct dynamic and flexible nature, anchored at the membrane via the Bcl-xL’s C-terminal helix and held together by the interaction of tBid’s BH3 domain with the Bcl-xL’s hydrophobic groove.

### The Bcl-xL/tBid complex at the membrane is a dimer with both structured and flexible regions

We created a series of active spin-labeled single cysteine variants of mouse cBid and human Bcl-xL (see Fig. 1 and Figs. S1, S2) for electron paramagnetic resonance (EPR) analysis. The chosen positions (*n*^Bid^ and *n*^Bcl-xL^) are depicted on the available structural models of water-soluble mouse Bid and human Bcl-xL in Fig. 1. Bcl-xL and cBid have different propensities to interact with the membrane: Bcl-xL partitions between aqueous and membrane environments (see for example [21, 32, 33]), while cBid preferentially stays in solution [34]. Notably, if tBid is purified alone, it readily interacts with the membrane [35]. This characteristic partitioning is found also for the spin-labeled variants of cBid and Bcl-xL when incubated alone in presence of LUVs (Fig. 2B). In addition, when the two proteins are mixed with LUVs, the partitioning of Bcl-xL is not affected, while cBid increases its tendency to separate into two fragments, with tBid being enriched at the membrane (Fig. 2B), in agreement with previous data on the wild type proteins [10, 36]. The effect exerted by Bcl-xL on the partitioning of tBid at the membrane provides a first indication of the sequestration of the BH3-only activator protein in an inhibitory membrane-bound complex. To structurally characterize this inhibitory complex, we measured a series of distance constraints between spin labels attached to cBid and Bcl-xL variants using Double Electron-Electron Resonance (DEER) spectroscopy (Fig. 2D,E). In the membrane-bound dimer, we found a defined distance between 82^Bid^, a label located few residues before the BH3 domain, and 110^Bcl-xL^, a label located in the hydrophobic groove (Fig. 2D). The obtained distance is perfectly in line with the BH3-into-groove interaction expected from the available water soluble crystal structure (PDB 4QVE, predicted distances in Fig. S7). This proves that the BH3-in-groove remains a key interaction between the two full-length proteins in the membrane-bound complex, confirming previous data obtained using full-length Bcl-xL and BH3-derived peptides (see for example [21, 37, 38]).

**Figure 2:**
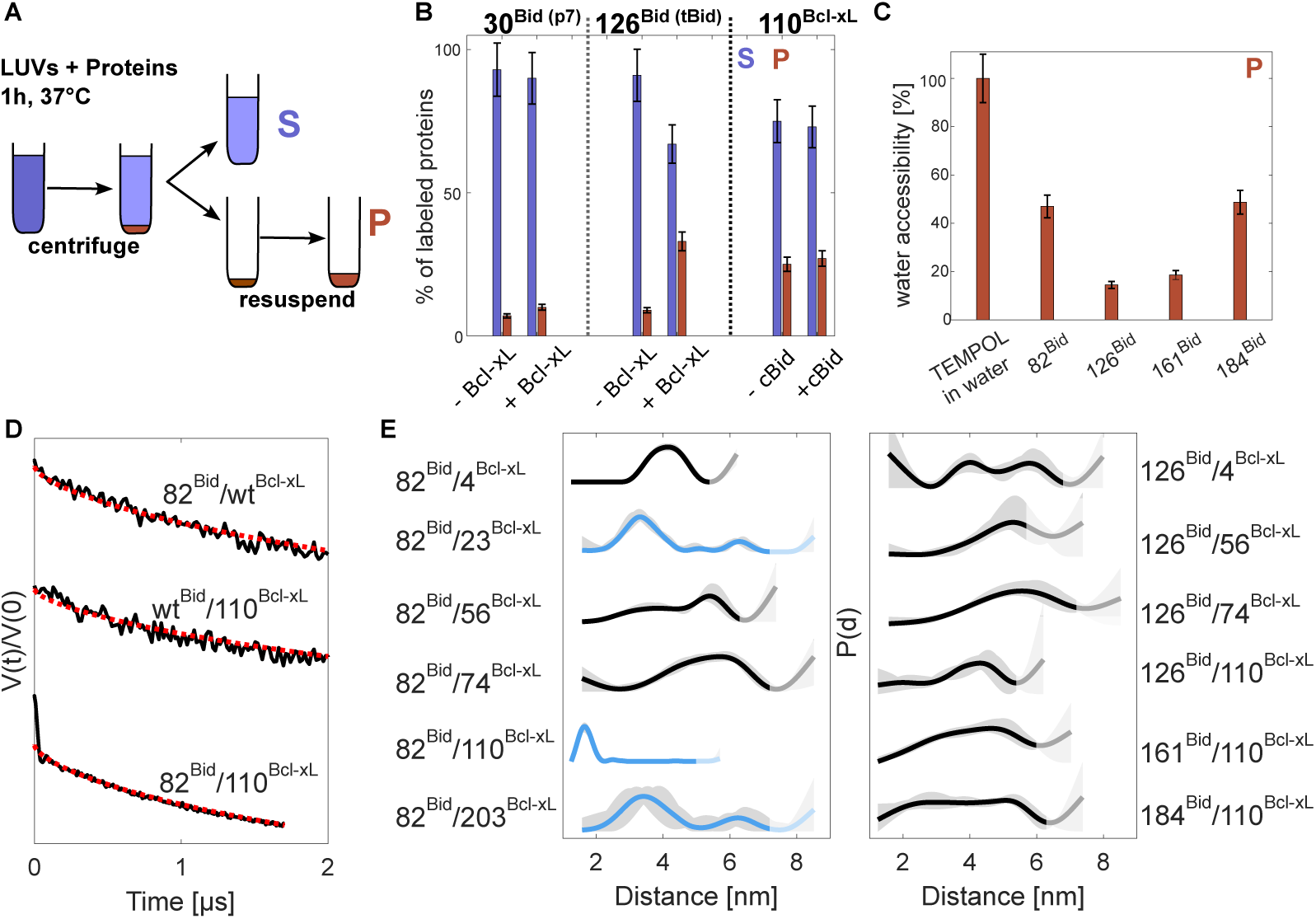
Experimental EPR data: cBid is sequestered by Bcl-xL at the membrane forming a heterodimer. **(A)** Scheme of the preparation of the aqueous (supernatant, S) and membrane (pellet, P) samples. **(B)** Fractions of p7 (30^Bid^), tBid (126^Bid^) and Bcl-xL (110^Bcl-xL^) partitioning in the supernatant and pellet samples either alone (-) or in presence (+) of the corresponding unlabeled protein partner (EPR spectra in Fig. S3). **(C)** ESEEM analysis of the water accessibility of tBid in complex with Bcl-xL at the membrane (P). The 100% reference sample is the water soluble radical TEMPOL. Primary data in Fig. S4. **(D)** Primary DEER traces of the pellet samples of 82^Bid^ incubated with unlabeled wild type Bcl-xL; unlabeled wild type cBid incubated with 110^Bcl-xL^; labeled 82^Bid^ and 110^Bcl-xL^ incubated together. The two upper traces (only one protein is labeled) are indistinguishable from a 2-dimensional background (red dotted), indicating the absence of specific homo-dimeric interactions. The bottom trace (one spin label in each protein) shows distinct dipolar oscillations due to the formation of a heterodimer. **(E)** Distance distributions (black, with shaded grey area representing the uncertainties) of different combinations of labeled cBid/Bcl-xL variants. Three distance constraints used for the initial MD model are in light blue. The shading of the distance distribution at long distances indicates the cutoff of the distance reliability. Primary data and additional DEER traces can be found in Figs. S5-8.

In addition, when each spin-labeled protein was mixed with its unlabeled partner, we could not detect specific Bcl-xL/Bcl-xL or cBid/cBid distances, proving that the inhibitory complex is a heterodimer and not a higher-order oligomer (Fig. 2D). Intriguingly, when probing the C-terminal region of cBid beyond residue 126, we could identify aspecific homo-interactions, indicating close proximity of the C-terminal regions of adjacent cBid proteins in the LUVs (see Fig. S6). All experimental Bcl-xL/cBid distances detected in the membrane-anchored heterodimer are presented in Fig. 2E. The first insight of the DEER analysis is that the BH3 domain of cBid (82^Bid^) has defined distances to residues in the globular region of Bcl-xL with full width at half maximum *<* 1.5 *nm* (compatible with the rotational freedom of the spin labels attached), indicating a tight binding of BH3 to Bcl-xL in the complex. Notably, broader distance distributions (*>* 2 *nm*) between the BH3 domain of cBid and two residues in the long *α*1*/*2 loop of Bcl-xL (56^Bcl-xL^, 74^Bcl-xL^) confirm that the loop of Bcl-xL remains flexible in the complex. The second insight is that the region of cBid following the BH3 domain is highly disordered, with distance distributions *>* 3 *nm* (Fig. 2E). In addition, based on the low accessibility towards deuterated water in the sample (detected via Electron Spin Echo Envelope Modulation, ESEEM), the C-terminal region of cBid is also found to be partially shielded from bulk water, suggesting its proximity to the membrane (Fig. 2C).

### Building the initial model of the Bcl-xL/tBid heterodimer

An initial model of the membrane-anchored human Bcl-xL/tBid complex was created with Modeller [39] following an integrative approach, combining existing partial experimental structures, homology modelling, and experimental observables [40]. The structures of the membrane-associated human tBid (PDB 2M5I [35], 64% sequence identity with the mouse Bid used for the DEER distance constraints, Fig. S9), a truncated aqueous complex of human Bcl-xL and a BH3 peptide from human Bid (PDB 4QVE [20]), and a model of the full-length membrane-embedded human Bcl-xL [21] were used as templates for the complex between tBid and Bcl-xL (see Fig. 3A). The template structures were positioned by aligning common structural regions. The EPR data on the water accessibility of cBid’s *α*4 − 8 helices (see Fig. 2) were used to re-orient the helices of tBid towards the membrane (see Fig. S10 and Methods).

**Figure 3:**
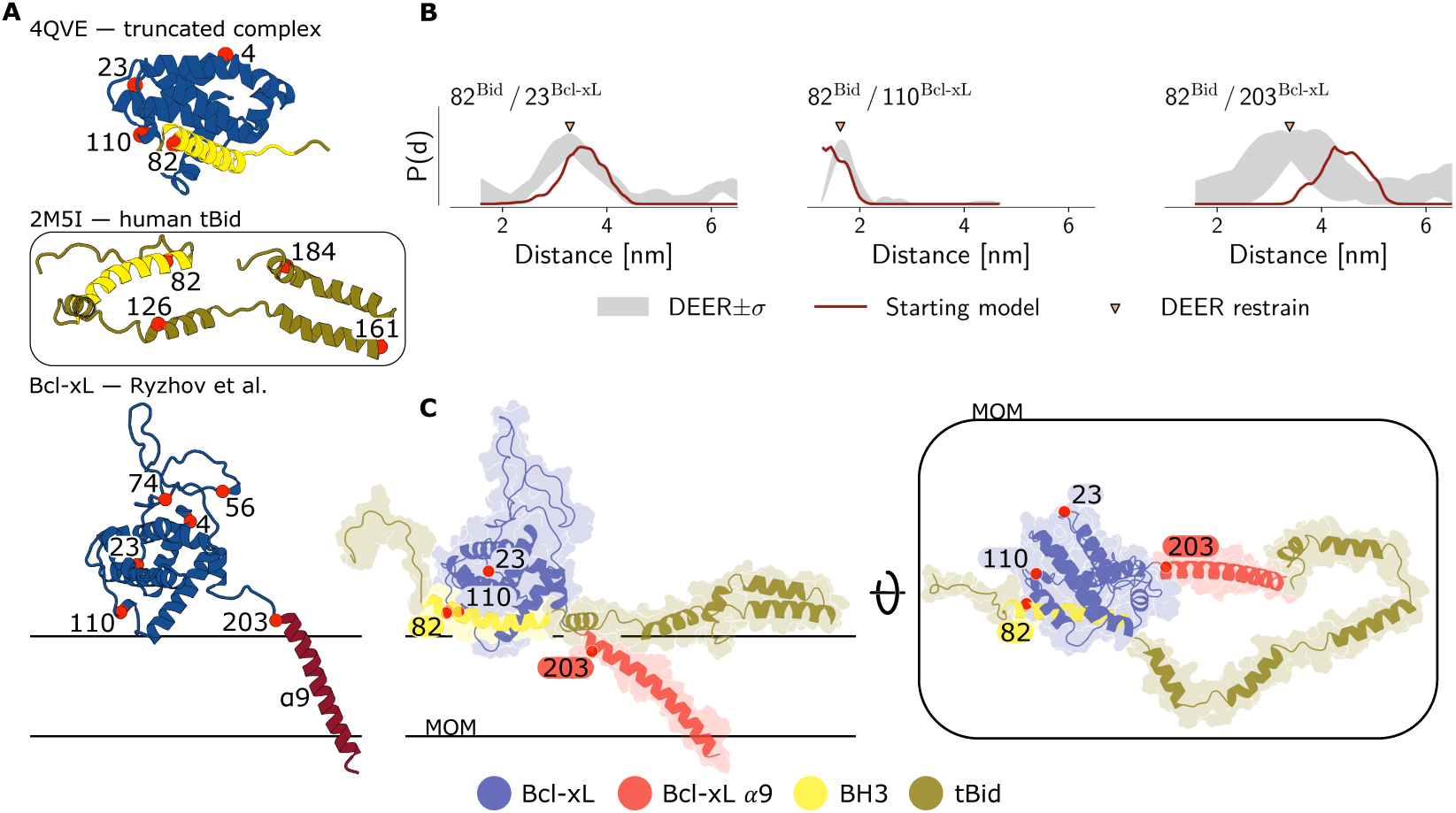
Initial model of the Bcl-xL/tBid complex at the membrane. **(A)** Reference structures used as basis for the homology modelling [20, 21, 35] **(B)** Three distance constraints (orange triangles) from Fig. 2E used to refine the starting model together with the ESEEM accessibility data from Fig. 2C. In red, the distance distributions calculated with DEER-PREdict [41] on the generated starting models. **(C)** Cartoon and surface representation of the starting model used for subsequent unbiased MD simulations. Label positions are indicated as red circles, the membrane (MOM) is indicated with black lines. The model was generated with Modeller [39].

Model generation was also guided by three experimental label-label distances between the BH3 domain of cBid and *α*1, *α*3, *α*8 of Bcl-xL (see Fig. 2D light blue lines). The resulting ten models showed a satisfactory initial agreement between the predicted and experimental label-label distances (see Fig. 3B). The model showing the best agreement to the EPR distances (see Fig. 3C and Methods) was used to run initial all-atom MD simulations (7 × 2 *µs*).

### A dynamic Bcl-xL/tBid inhibitory complex at the membrane

Capturing the full flexibility of the Bcl-xL/tBid complex proved difficult within initial all-atom simulations (see Fig. S11). To diversify the starting models of all-atom simulations and access a larger conformational landscape we re-seeded all-atom simulations (12 × 2 *µs*) from coarse grained simulations (9 × 7 *µs*) of the complex (see Fig. S12 and Methods).

The re-seeded all-atom ensemble is characterised by both structured and highly flexible regions in the complex (Fig. 4A,B,C), in line with the experimental EPR data (see Fig. 2E). Four independently moving domains could be identified within the complex (see Fig. S15D).

**Figure 4:**
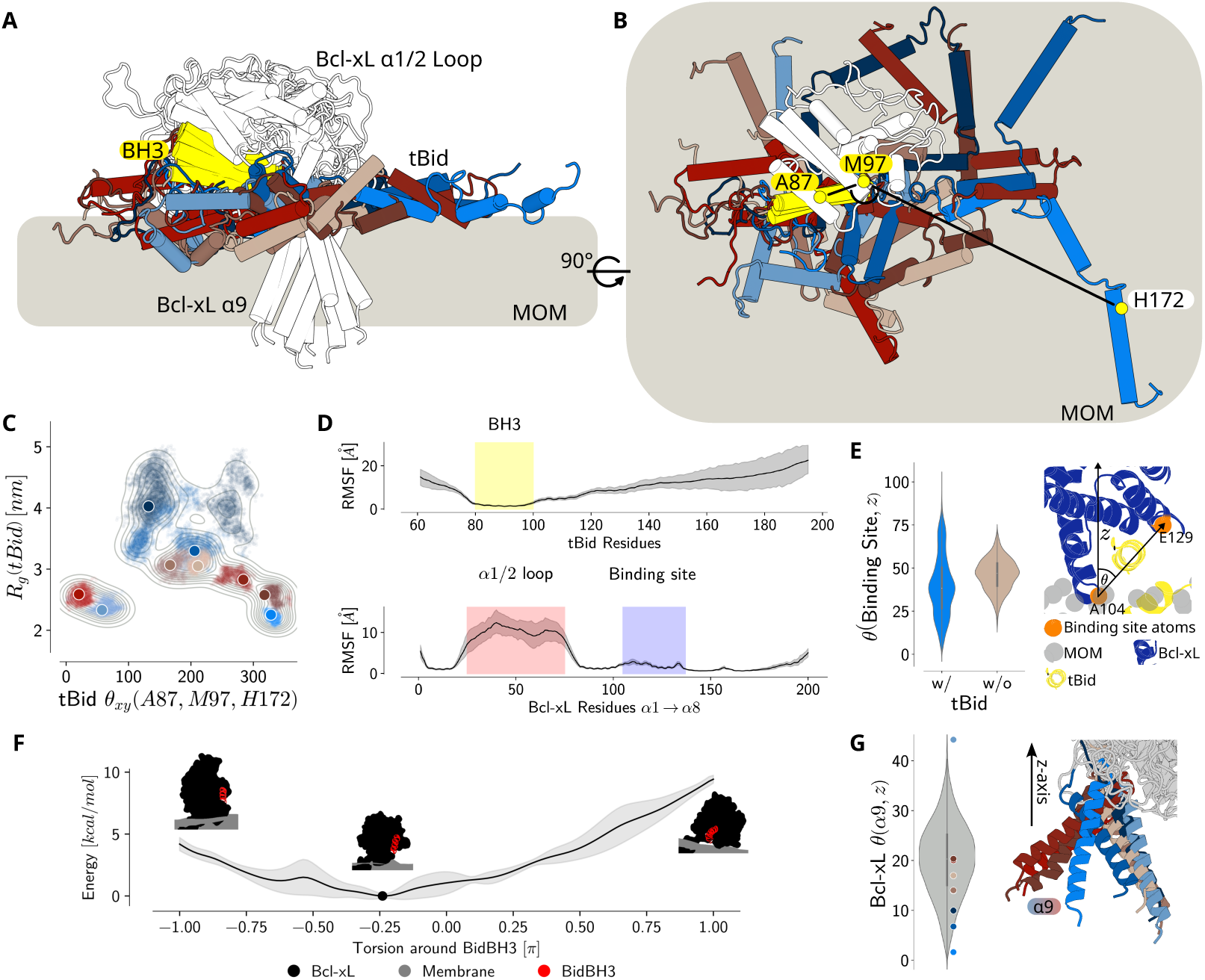
Structured and dynamic regions coexist in the Bcl-xL/tBid heterodimer. **(A)** Side view of the cluster representatives of the Bcl-xL/tBid complex structural ensemble embedded on a lipid membrane. The representative of tBid are colored in shades of red, brown, blue (cartoon representations) with the BH3 domain shown in yellow. The cluster representatives of Bcl-xL are shown in white. **(B)** Top view of the cluster representatives to highlight that tBid’s helices *α*4 − 8 diffuse freely on the membrane. For clarity, only one cluster representatives of Bcl-xL is shown in white. **(C)** Scatter plot of tBid’s radius of gyration (*R_g_*) and angular spread (Θ*_xy_*, angle-defining atoms shown as yellow spheres in (B)). Cluster representatives indicated as larger colored dots. **(D)** Root mean square fluctuations (RMSF) of tBid and Bcl-xL. Structures were aligned on the *α*5 and *α*6 helical C*α*-atoms of Bcl-xL. Specific regions of the sequences are highlighted. **(E)** The hydrophobic groove of Bcl-xL is tilted away from the membrane when tBid is bound. The binding site is shown as cartoon (Bcl-xL: blue; tBid: yellow). The plot shows the comparison of the angular spread of the binding site of Bcl-xL alone or in complex with tBid (vector connecting A104 and E129, orange spheres) with respect to the membrane normal (z). **(F)** In the complex, the BH3 is located between the hydrophobic groove and the membrane headgroup region as shown by the free energy potential of the torsion between Bcl-xL’s globular domain and the membrane lateral over the BH3 helix. **(G)** Angular spread of the *α*9 of Bcl-xL with respect to the membrane normal (z) indicated as dots in the violin plot. Cluster representatives of the *α*9 of Bcl-xL are shown in shades of red, brown and blue (corresponding to the representatives of tBid in panel A), aligned on the globular domain of Bcl-xL. Additional information can be found in Figs. S11-16.

The amphipathic alpha helix of tBid containing the BH3 domain remains tightly bound to Bcl-xL’s globular domain via the BH3-in-groove interaction (see Fig. 4A,B,D and Tab. S1), forming the largest independently moving domain (see Fig. S15D). In contrast to the published aqueous structure [20], the hydrophilic surface of Bid’s BH3 domain is not facing the bulk water; instead, it is oriented towards the membrane headgroup region (see Fig. 4E). In the re-seeded MD trajectories, the average distance between the center of Bid’s BH3 domain (residue E96 pointing towards the membrane) and the lipid phosphates is 0.59 ±0.22 *nm*. Notably, the groove of Bcl-xL slightly moves away from the membrane upon tBid binding (see Fig. 4E) based on MD simulations. To further corroborate the binding of tBid in between the groove and the membrane headgroups, we sampled the orientation of the globular domain of the complex relative to the membrane through enhanced sampling (see Fig. S13) [42, 43]. The obtained free energy surface (FES) supports the proposed membrane-facing orientation as the most likely one (see Fig. 4F), which was also corroborated by FES estimation of the re-seeded all-atom simulations (Fig. S13E).

The second largest domain consists of the tBid C-terminal helices freely diffusing on the membrane (see Fig. S15C-E), adopting both compact and extended conformations (see Fig. 4 B,C). These C-terminal helices are able to form transient contacts with the globular domain of Bcl-xL in either a clockwise or counter-clockwise fashion (see Fig. 4C Tab. S1).

The complex is membrane-anchored via the *α*9 (see Fig. S15C,D) of Bcl-xL, which moves independently from the largest domains (see Fig. S16). This TMD retains a tilted orientation in relation to the membrane normal (see Fig. 4G), as shown in previous studies [21]. The globular domain of Bcl-xL contacts the membrane predominantly via the *α*2 helix, the C-terminal tip of the *α*1 helix, the *α*8 helix, and transiently also via the *α*4*/*5 loop (see Fig. S15C). The remaining smaller domain is the flexible *α*1*/*2 loop of Bcl-xL (see Fig. 4D and S15D), whose flexibility increases when Bcl-xL binds tBid (see Fig. S16). Binding of tBid also modifies the dynamics in the binding site of Bcl-xL, increasing its local flexibility (see Fig. S16). To summarize, Bcl-xL sequesters tBid at the membrane, locking it in between the canonical hydrophobic groove and the membrane’s headgroups. The resulting complex is a highly flexible heterodimer, tethered via the Bcl-xL’s *α*9 helix to the membrane, with the C-terminal region of tBid freely diffusing on the membrane.

### *In vitro* and *in organelle* experiments validate the Bcl-xL/tBid complex

The atomistic Bcl-xL/tBid model was validated through independent experiments *in vitro* and *in organelle*.

First, hydrogen deuterium exchange mass spectrometry (HDX-MS) was used to probe the solvent accessibility of the complex in LUVs, highlighting fold dynamics, membrane-protein and protein-protein interactions [44]. Deuterium exchange patterns within Bcl-xL from the water soluble to the membrane-embedded state confirm the membrane-embedding of the *α*9 helix in Bcl-xL alone and in presence of cBid, with decreased relative deuteration in the latter case (see Fig. S17). Simulations on the ensemble Bcl-xL/tBid dimeric model show excellent correlation with the experimental deuteration patterns of Bcl-xL in the membrane fraction in presence of cBid (0.76 ≤ *R*^2^ ≤ 0.92) (Fig. 5B), which validated structure, dynamics and topology of the anti-apoptotic protein in the complex. The experimental HDX pattern of cBid indicates an increase in the protein’s deuteration and flexibility going from the water-soluble form to the complex with Bcl-xL at the membrane (see Fig. S19), corroborating the high flexibility observed in the C-terminal region of tBid in the dimeric ensemble (see Fig. 4C). The simulated HDX data shows a reasonable correlation for the human tBid (0.15 ≤ *R*^2^ ≤ 0.38) in the heterodimeric ensemble with the experimental patterns obtained with the mouse tBid (see Fig. 5C). The minor discrepancies observed might be due to small variations in the peptide sequences of mouse and human tBid (Fig. S9) and the residual presence of water soluble cBid in the sample (see Fig. 4C and Fig. S19).

In a second step the dynamic model of the inhibitory complex was further validated by comparison of the label-label DEER distance distributions predicted on the re-seeded MD trajectories with the experimental data (see Fig. 5A). The agreement between simulations and *in vitro* distance constraints is good and lies within the expected root mean square deviations of the rotamer library approach used [45]. The DEER validation therefore supports the binding mode between Bcl-xLand tBid obtained via MD simulations (see Fig. 5D). Lastly, to corroborate the physiological relevance of this dynamic inhibitory complex, we investigated four key distances in isolated mitochondria from mouse liver. The inhibitory Bcl-xL/tBid complex formed in mitochondria showed the same distance constraints found *in vitro* between the BH3 domain and Bcl-xL’s globular domain, and the broad distance distribution between tBid’s C-terminal helices and Bcl-xL (see Fig. 5E). The data in mitochondria confirm that tBid binds Bcl-xL’s hydrophobic groove via its BH3 domain and that the following helices have a high degree of flexibility in a physiologically relevant environment.

**Figure 5:**
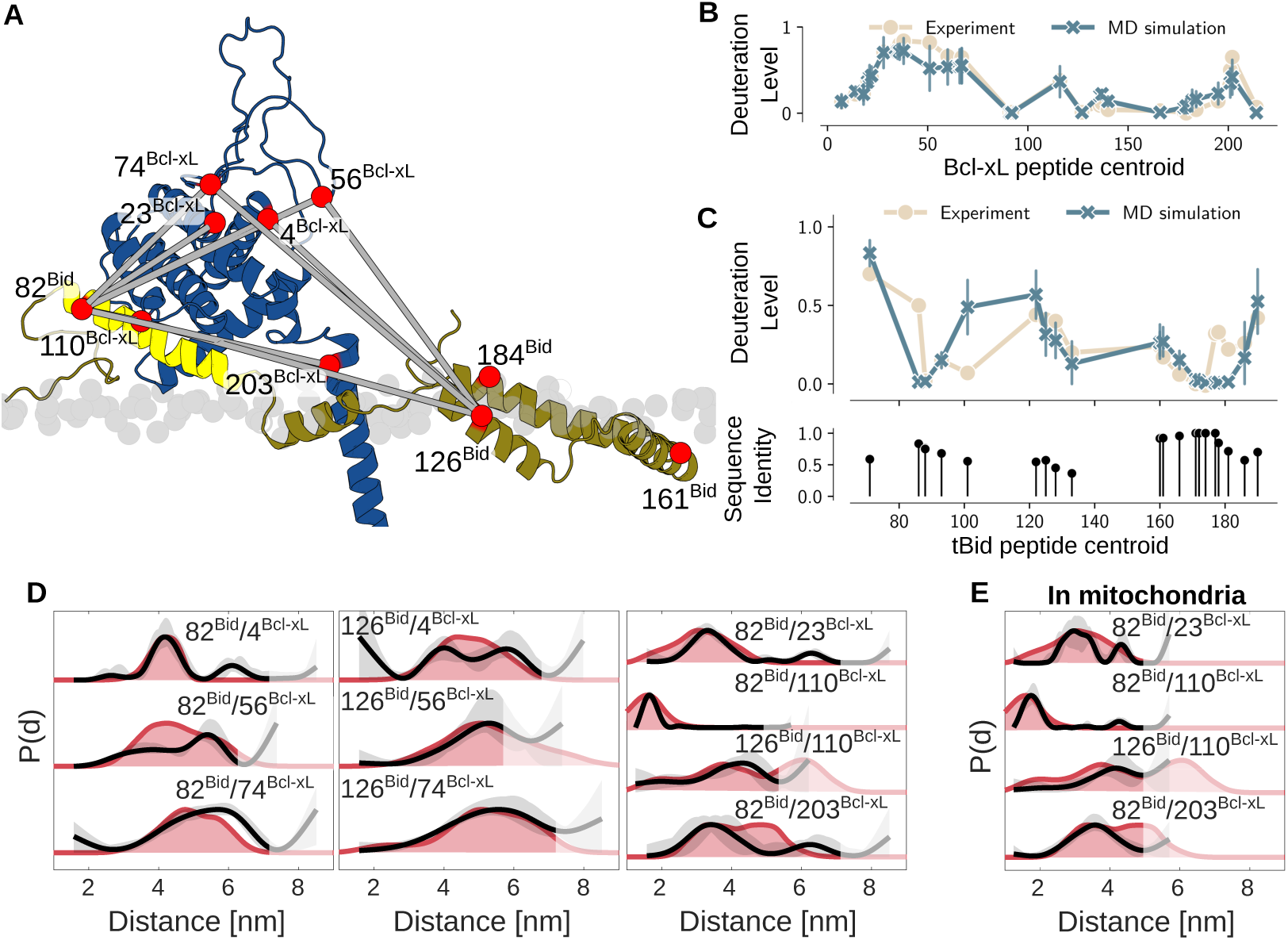
Experimental validation of the structural ensemble. **(A)** Representative cartoon model of the tBid/Bcl-xL complex (blue: Bcl-xL, yellow: BH3 domain of tBid, dark yellow: tBid, light grey: lipid phosphates) with indicated label positions (red spheres). The experimental distances chosen to validate the structural model (grey lines) are unambiguously assigned to Bcl-xL/tBid interactions. The experimental broad distance distributions towards positions 161 and 184 in tBid (Fig. 2E) were omitted from the validation because they also contain aspecific tBid/tBid interactions (Fig. S6). **(B)** Experimental (beige, pellet fraction after incubation of cBid with Bcl-xL and LUVs) versus simulated (blue, Bcl-xL/tBid complex) deuterium uptake of Bcl-xL peptides (additional data in Figs. S17,18). **(C)** Experimental (beige, pellet fraction after incubation of cBid with Bcl-xL and LUVs) versus simulated (blue, Bcl-xL/tBid complex) deuterium uptake of tBid peptides (additional data in Figs. S18,19). The sequence identity between human (MD simulation) and mouse (experiment) tBid for the detected peptides is shown. **(D)** Experimental DEER distance distributions (black with grey area representing the uncertainty and the shaded region indicating the cutoff for reliable distances according to [46]) compared to those calculated on the MD simulation trajectories using DEER-PREdict (red). Additional distance distributions, technical and biological repeats, primary data, and analysis can be found in Figs. S7,8. **(E)** *In organelle* distance distributions (color code as in panel D) compared to the corresponding calculated distance distributions. Additional information can be found in Figs. S20,21,23.

## Discussion

Resolving the interaction mode of MOM-associated Bcl-2 inhibitory complexes is crucial for understanding the regulation of apoptosis and for developing potent apoptotic modulators. Here, using an integrative modeling approach that combines experimental data with molecular simulations, we describe the structure, interaction interfaces, and dynamics of the inhibitory complex between Bcl-xL and tBid at the mitochondrial outer membrane.

Bcl-xL has been reported to shuttle between the cytosol and the membrane [33, 47]. Its membrane-anchored form is responsible for sequestering tBid in the indirect MOMP inhibition mode. By competing with p7 [34], Bcl-xL shifts tBid’s equilibrium towards the membrane-bound form and finally sequesters it. Our analysis reveals a structural dichotomy in the membrane-bound Bcl-xL/tBid complex. While tBid’s BH3 domain is firmly locked between Bcl-xL’s hydrophobic groove and the membrane bilayer, impeding any interaction with effector proteins, its C-terminal region remains flexible and diffuses freely across the membrane surface. This mobile C-terminus, together with the TMD of Bcl-xL, forms the complex’s extensive membrane-interaction surface (see Fig. 4D and Fig. S15C). These flexible regions may even permit loose interactions between adjacent heterodimers. Whether such crowding contributes to the sequestration mechanism or regulates interactions with other BH3-only proteins remains an open question. The flexibility of the C-terminal helices of tBid in the inhibitory complex is reminiscent of the flexibility induced on Bak by the interaction with the anti-apoptotic protein MCL-1 [48], which is intriguing for mechanistic implications. Notably, DEER cannot distinguish between partially unfolded and dynamic helices, but we have evidence by ESEEM that the C-terminal region of tBid remains at the membrane, suggesting the presence of alpha helices as in the NMR structure of isolated tBid in nanodiscs [35] and we have no hints of unfolding events during the extended unbiased MD simulations. This inhibitory mechanism highlights the crucial role of the MOM in shaping the binding interface, a feature necessarily absent in previous studies using truncated, water-soluble proteins. Although we did not simulate the initial binding event, our model reveals its structural consequence: the globular domain of membrane-anchored Bcl-xL repositions itself away from the membrane upon tBid binding. This movement exposes the hydrophobic groove, effectively locking the BH3 helix in between the groove and the membrane headgroup region, in stark contrast to the more accessible binding site of truncated Bcl-xL variants in solution [20]. Notably, this observation aligns with the proposed “cryptic” nature of Bcl-xL’s binding pocket, which is thought to require unfolding for ligand access [49–51]. Indeed, the *α*3 helix of Bcl-xL, which was unfolded in our starting model [21], remains so throughout our simulations. We find that Bcl-xL sequesters pro-apoptotic tBid exclusively at the membrane, as any interaction in the cytosol is sterically prohibited. This prevention occurs because the BH3 domain of cBid is occluded by its p7 fragment, while the corresponding binding groove on Bcl-xL is blocked by its own *α*9 helix. The membrane is therefore an essential component of the inhibitory complex, which recruits the protein partners, shapes the BH3 binding site and allows the flexibility of the C-terminal helices of tBid. Indeed, AlphaFold2 [52] fails to correctly predict the topology and assembly of the inhibitory complex due to the lack of the membrane environment folding tBid’s C-terminal helices around Bcl-xL’s transmembrane helix (see Fig. S22).

The mechanism of tBid inhibition fundamentally differs from how Bcl-xL inhibits Bim, a BH3-only protein that, unlike Bid, contains a C-terminal transmembrane domain (TMD) [53]. Bcl-xL was found to neutralize Bim via a “double-bolt” lock mechanism, engaging both the canonical BH3-in-groove interface and a direct TMD-TMD interaction. This robust, dual-mode sequestration might contribute to cellular resistance against BH3-mimetic drugs that target only the binding groove. In addition, Bcl-xL was found to inhibit apoptosis as a higher-order complex that binds multiple BH3 proteins [32]. The existence of distinct inhibitory strategies will require development of therapeutic approaches tailored to specific protein-protein and protein-membrane interfaces [47, 53–56]. Elucidating the structural basis for the diverse inhibitory mechanisms within the Bcl-2 family is therefore paramount for designing next-generation therapies.

## Materials & Methods

### Protein expression, purification, and labeling

His-tagged mouse Bid (pET23d [57]), human Bax (pTYB1 [58]), and human Bcl-xL (pTYB1 [59]) were prepared as described previously [36].

In short, all plasmids except for pTYB1-Bcl-xLC23C203 (S23CC151SS203C, ordered from GenScript) were cloned by site-specific mutagenesis, which was confirmed by DNA sequencing. All proteins were expressed in *E. coli* and the proteins were purified from the soluble fraction of the bacterial lysate.

Bid was purified using Ni-NTA and washed with buffer in three steps (imidazol: 0 mM, 10 mM, 25mM) before being eluted with 250 mM imidazole. For cleavage, Bid was mixed 1:1 with cleavage buffer and cleaved with Caspase 8. Afterwards, cBid was purified with Ni-NTA.

cBid was spin labeled with MTSL using a standard protocol: 10-20 µM protein concentrations were incubated with 1 mM DTT for 1 h. DTT was removed in a desalting column, and the label was immediately added (10x excess) from a concentrated stock solution in DMSO. The solution was incubated at 8°C overnight and the excess label was removed using a desalting column.

Bax and Bcl-xL were purified using a chitin resin and washed thoroughly before cleaving the intein tag off overnight by incubation with DTT. After elution, Bcl-xL was dialyzed in one step 1:10 against 20 mM Tris (pH 8.0) and Bax was dialyzed in five steps against 8 mM Tris (pH 8.0). Both proteins were further purified using anion exchange columns. For Bcl-xL 1 mM DTT was added to the buffers forloading, washing, and elution.

The pooled fractions of Bcl-xL were purified in a third step using a Superdex 200 10/300. If the protein had to be labeled, 10 *µ*M TCEP was added to the buffers.

MTSL-labeling of Bax and Bcl-xL was performed with protein concentrations between 10 and 20 *µ*M and 5-10x excess of MTSL at 8°C overnight. The excess label was then removed by washing in centrifugal filters.

All proteins were tested in pore formation assays (Fig. S2) before further experimental use.

### LUVs preparation

Large unilamellar vesicles (LUVs) for EPR, HDX and pore formations assays were prepared as described before [36] with a composition mimicking the MOM (46% egg L-*α*-phosphatidyl choline (PC), 25% egg L-*α* phosphatidyl ethanolamine (PE), 11% bovine liver L-*α*-phosphatidyl inositol (PI), 10% 18:1 phosphatidyl serine (PS) and 8% cardiolipin (CL) (% w/w)). In short, the dried lipid mixture was resuspended either in storage buffer (EPR, HDX) or in calcein solutions (80 mM, pore formation assays) and extruded through a membrane with 400 nm pores. As control, a lipid mixture corresponding to the exact composition used in the MD simulations was prepared the same way with 80% 1-palmitoyl-2-oleoyl-sn-*glycero*-3-phosphocholine (POPC) and 20% CL.

### Pore formation assays

The pore formation assays were performed as decribed before [36]. In short, the calcein was trapped in the LUVs at a high, self-quenching concentration, leading to a low fluorescence intensity. Upon pore formation in the membrane, calcein is diluted in the surrounding buffer leading to increasing fluorescence intensity. All proteins were used at 25 nM final concentration.

### Continuous wave EPR

Room temperature CW EPR experiments were performed on a Bruker E500 X-band equipped with a super high Q cavity ER 4122 SHQ with a power of 2 mW and a modulation amplitude of 0.1 or 0.15 mT. Samples were measured in glass capillaries (Blaubrand). Spin counting was performed by comparison of the second integral of the EPR spectrum with the second integral of a reference spectrum of a 100 *µ*M TEMPOL solution in water.

### Pulsed EPR measurements

All pulsed EPR experiments were performed at 50 K on a Bruker Q-band Elexsys E580 spectrometer equipped with a 150 W TWT amplifier, a Bruker Spinjet-AWG, and a homemade probe head accommodating 3 mm o.d. tubes. The temperature was stabilized using a cryo-free system from Cryogenics Limited.

The dead-time free 4-pulse DEER sequence with Gaussian pulses [60] was used with 36 ns Gaussian pulses. The pump position was chosen at the maximum of the nitroxide spectrum and the observer position at −90 MHz. The upper limit of accurate mean distances is indicated with the shaded region in all DEER distance distributions shown, and it was determined by the following formula [46]:

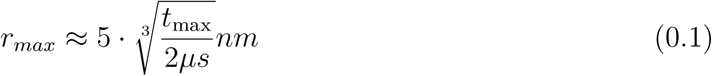

The parameters used are summarized in Table S2.

For ESEEM we used the 3-pulse sequence with rectangular 12 ns pulses and 4-step phase cycling to eliminate running echoes. *τ* was optimized in 2D 3-pulse ESEEM experiments to maximize the deuterium signals. The parameters used are summarized in Table S3.

Solution samples were prepared by mixing 20 *µ*l protein sample with 20 *µ*l d8-glycerol. Membrane samples were prepared by incubating the protein mixtures with LUVs for 1 h at 37°C and by separating the pellet from the supernatant by centrifugation. The pellet was resuspended in buffer prepared with deuterated water and mixed with 10% d8-glycerol to the final sample volume of 40 µl. The samples were shock frozen in 3 mm o.d. quartz tubes (Aachener Quarzglas) and stored at −80°C.

DEER data were analyzed with DeerAnalysis [61] 2022 using Tikhonov regularization. ESEEM data were analyzed within the Bruker Xepr software by fitting the trace with a biexponential decay and dividing the data by the fit for baseline correction. After application of a Hamming function and zero filling, the data were Fourier transformed to obtain the intensity of the deuterium signals used as descriptors of the water accessibility of the spin-labeled side chain.

### Hydrogen/deuterium exchange mass spectrometry

HDX-MS experiments were performed at the UniGe Protein Biochemistry Platform (University of Geneva, Switzerland) following a well-established protocol with minimal modifications [62]. The following conditions were tested and compared: i) Bcl-xL; ii) Bcl-xL + LUVs; iii) Bcl-xL + LUVs + cBid and iv) cBid. For Bxl-xL and cBid, a first concentrated mix was prepared and preincubated for 5 min at the reaction temperature before initiating deuteration reaction. For reactions containing LUVs (conditions ii and iii), proteins (340 pmol) were pre-incubated with LUVs for 1h at 37 °C. The mix was then centrifuged at 20’000 ×g for 30 min at 22 °C and the supernatant containing proteins unbound to lipids was removed. The pelleted material was used for deuterium labeling.

Deuterium exchange reaction was initiated by adding deuterated buffer at the defined temperature to a final reaction volume of 50 microliters. Reactions were carried-out for defined times and terminated by the sequential addition of ice-cold quench buffer. Samples were immediately frozen in liquid nitrogen and stored at −80 °C for up to two weeks.

To quantify deuterium uptake into the protein, samples were thawed and injected in a UPLC system immersed in ice with 0.1 % FA as liquid phase. The protein was digested via an immobilized Nepenthesin-2/Pepsin mixed column (AffiPro AP-PC-006), and peptides were collected onto a Nucleodur 300-5 C18 pre-column filter (Macherey-Nagel). The trap was subsequently eluted, and peptides separated with a C18, 175 Å, 1.9 *µ*m particle size Thermo Hypersil Gold Vanquish 100 x 2.1 mm column over a gradient of 8 – 30 % buffer C over 20 min at 150 *µ*l/min (Buffer B: 0.1% formic acid; buffer C: 100% acetonitrile). Mass spectra were acquired on an Orbitrap Velos Pro (Thermo), for ions from 400 to 2200 m/z using an electrospray ionization source operated at 300 °C, 5 kV of ion spray voltage. Peptides were identified by data-dependent acquisition of a non-deuterated sample after MS/MS and data were analyzed by Mascot 2.6 using a database composed of purified proteins and known contaminants. Precursor mass tolerance was set to 10 ppm and fragment mass tolerance to 0.6 Da. Protein digestion was set as nonspecific. All peptides analysed are shown in HDX Tables. Deuterium incorporation levels were quantified using HD examiner version 3.4.2 software (Sierra Analytics), and quality of every peptide was checked manually. Results are presented as percentage of maximal deuteration compared to theoretical maximal deuteration. Criteria to define changes in deuteration level between two states as significant are indicated in the tables.

### Mitochondria purification

Crude mitochondria were isolated from mouse liver as described in literature [63]. In short, fresh mouse livers with a total of approximately 3.2 g were cut in small pieces on ice and homogenized in sucrose buffer using a glass dounce homogenizer. The homogenate was centrifuged at 600×*g* for 10 minutes at 4°C to remove any cell debris. Subsequentially, the supernatant was centrifuged at 7000×*g* for 10 minutes at 4°C. Approximately 1 mL of crude mitochondria suspension in sucrose buffer was then obtained and kept on ice. Mitochondrial proteins were considered as an index of mitochondrial concentration, and they were determined using a Bradford assay. The final protein concentration was around 90 mg/mL. For DEER samples mitochondria (final amount: ∼ 1 mg mitochondrial protein) were mixed with proteins (final amount: ∼ 0.5 nmol) and incubated for 5 min at 37°C. Then they were centrifuged for 3 min at 7000×g and 4°C. The supernatant was removed and the mitochondria were resuspended in 30 *µ*l deuterated Threalose buffer (200 mM Trehalose, 10 mM HEPES, 10 mM EGTA, 10 mM EDTA) to the final sample volume of 40 µl. The samples were shock frozen in 3 mm o.d. quartz tubes (Aachener Quarzglas).

### Model generation

Hetero-complex models were based on a model of membrane-associated Bcl-xL by Ryzhov et al. [21], PDBid 2m5i [35] and PDBid 4qve [20]. As such the model comprised human Bcl-xL and tBid. Models of the hetero-complex were generated with custom modelling classes within the Modeller python software package [40]. Input reference PDB structures for Modeller [40] (see Fig. S10) were prepared as follows:

1. The BH3 helix and N-terminal loop was removed from all models within PDB 2m5i [35].
2. From the apo Bcl-xL model the loop connecting helix *α*8 to *α*9 was removed.
3. Modified apo Bcl-xL model was aligned to the Bcl-xL chain in PDB 4qve [20].
4. Modified 2m5i structures were positioned such that the N-terminal is close to the C-terminal of the BidBH3 in 4qve and parallel to the membrane plane of the modified apo Bcl-xL model, such that the membrane interacting residues point towards the membrane [21, 35].

Alignments and placements were made with PyMOL [64]. Models were generated using the LoopModel class of Modeller with the following additional restraints: (i) Bcl-xL was rigid body restrained except in the *α*1*/*2 loop and after helix *α*8, (ii) Alpha helices *α*9 of Bcl-xL, *α*4, and *α*5 of tBid were secondary structure restrained, (iii) the centre of mass distance between DEER residue pairs were restrained towards the global maximum of the DEER probability distributions with the Gaussian restraint form in Modeller and a standard deviation of 0.5 Å, and (iv) the z positions of atoms in the tBid N-terminal loop were lower bound restraint to be above the membrane plane implied by the modified apo Bcl-xL model. Only the beginning of the Bcl-xL *α*1*/*2 loop (residues 21 to 27) and tBid N-terminal loop were selected during loop remodelling. Initial modelling used the autoscheduler with maximum 600 conjugate gradient iterations, no MD refinement and 14 optimization repeats. Loop modeling used the fast MD refinement and ten optimization repeats. Models were discarded if their molpdf score was above 2 × 10^10^. Generated output models were re-scored based on their DOPEHR score and a DEER score as per the following:

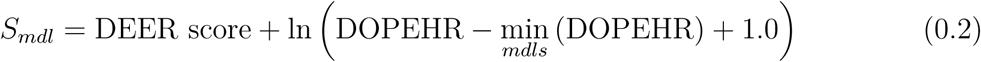

The DEER score of a model is computed as the sum of all Wasserstein distances between normalised^1^ experimental DEER distance distributions and their corresponding estimated distributions in the model. DEER distributions of distances in the models were estimated using DEER-PREdict [41, 65]. Briefly, DEER-PREdict places a rotamer library of the DEER label onto the distance partners, removing clashing rotamers and calculates the distance distribution between the paired electrons. Models were protonated at pH 7.5 using pdb2pqr [66] and propka [67] giving a total charge of −9*e* to the hetero-complex.

### All-atom molecular dynamics simulations

All simulations were run with GROMACS 2023 [68] and plumed 2.9.0 [43]. For water the modified CHARMM36m water parameters were used and for lipids, ions, and protein the CHARMM36m forcefield was used [69]. A breakdown of changing simulation settings throughout the equilibration and production simulations can be found in Tab. S4. In NVT and NPT simulations the V-rescale thermostat was employed at a temperature of 310.15 *K* with a time constant of 1.0 *ps*, separately coupling membrane, protein and solvent [70]. NPT simulations used a semi-isotropic C-rescale barostat [71] at 1 *atm* with a compressibility of 4.5 × 10^−5^ *bar*^−1^ and a time constant of 5.0 *ps*. Neighbour lists were build with the Verlet cutoff scheme. Lennard-Jones interactions were treated with the Cut-off scheme using force-switching at 1.2 *nm* and switch radius of 1.0 *nm*. Coulomb interactions were calculated using particle-mesh-Ewald [72] with a cutoff 1.2 *nm*. Hydrogen bonds were LINCS [73] constrained and center of mass motion removed every 100 steps on the solvent and solute groups separately. Frames were written to trajectory at a frequency of 10 *ns*^−1^.

### Initial Bcl-xL/tBid all-atom simulations

The protonated highest scoring Bcl-xL/tBid (holo) model was solvated with CHARMM-GUI [74] in a hexagonal box containing 576 (80 %) POPC and 144 (20 %) 9,10 and 12,13 unsaturated cardiolipin (TLCL) (charge: −2*e*). This membrane composition was tested for Bcl-xL activity and complex formation in the experimental setup and showed comparable distance distributions to other conditions (Fig. S20). Lipids were equally distributed across the leaflets and the system solvated in TIP3P water [69]. The box was charge neutralised with sodium chloride and sodium chloride atoms added to a 150 *mM* concentration. Within the box the protein was positioned such that tBid is slightly above the membrane plane to allow for proper lipid placement by CHARMM-GUI.

After energy minimization and equilibration tBid was embedded into the membrane using adaptive biasing MD [75]. Embedding was described by the minimum value of the difference in coordination of each hydrophobic residue in tBid with lipid phosphates and carbons. The secondary structure of tBid was restrained by the sum of all backbone hydrogen bonds in alpha helices. Once reasonable embedding was achieved, adaptive biasing MD simulations were stopped. Embedding was followed by a 15 *ns* relaxation in which only the secondary structure of tBid and the lipid phosphate positions were restrained. Phosphates were restrained towards the mean value in the z-dimension of the initial frame shifted towards tBid by 0.5 Å and secondary structure restrained as before (Tab. S4). For production simulations the equilibrated and embedded system was copied seven times to run seven independent replicates. First 60 *ns* of unrestrained MD were run with new initial velocities and discarded as equilibration, to which followed seven production simulations. Each production replicate has a duration of 2 *µs* (Fig. S11).

### All-atom Bcl-xL simulations

Bcl-xL only (apo) systems were generated with CHARMM-GUI from the Bcl-xL apo model [21]. The protein was embedded in 20 % TLCL and 80 % POPC at 150 *mM* NaCl and neutralising ions. Four independent replicates were equilibrated from the generated system (Tab. S4), followed by 200 *ns* of production simulation.

### Coarse grained Bcl-xL/tBid simulations

Coarse grained simulations were initialized from cluster representative structures identified in the initial all-atom simulations (Fig. S11). The simulation boxes for the nine cluster representatives were generated with martinize2 and insane [76, 77] within a membrane of 80%*/*20% POPC/TLCL2 solvated with MARTINI water, neutralizing ions (Na+) and 150 *mM* NaCl salt. Boxes were parameterized with the intrinsically disordered protein version of MARTINI3 [78–80]. An elastic network independently generated in each replicate was applied to the protein with a force constant of *κ* = 700 *kJ/mol*, excluding the *α*1*/*2 loop of Bcl-xL and loops between helices of tBid. Simulations were run with GROMACS 2023 in the NPT ensembles after equilibration (Tab. S4). Temperature was regulated with the V-rescale thermostat [70] at 310.15 *K* and a time constant of 1.0 *ps*. Pressure was coupled semi-isotropically via the C-rescale barostat [71] at 1 *atm* with a compressibility of 3 × 10^−4^ and a time constant of 4.0 *ps*. Neighbour lists were updated every 20 steps and build with the Verlet cutoff scheme under a buffer tolerance of −1 [81]. Lennard-Jones interactions were treated with the Cut-off scheme using potential shifting with a switch radius of 1.1 *nm*. Coulomb interactions were calculated using reaction-field with a radius of 1.1 *nm*. No constraints were applied and frames written 1 *ns*^−1^. Each simulation had a length of 7 *µs* (9 × 7 *µs*).

### Re-seeded Bcl-xL/tBid all-atom simulations

From the coarse-grained simulations twelve cluster representatives (Fig. S12) were extracted via ELViM dimensionality reduction and gaussian mixture model clustering [82]. Protein and membrane of extracted structures were backmapped into CHARMM36m parameterized all-atom representations using the Multiscale simulation tool [83]. Consecutively backmapped structures were re-solvated (see all-atom molecular dynamics simulations). Replicates were equilibrated under restraint on the *α*-helical hydrogen-bonds between backbone atoms of Bcl-xL and tBid to retain secondary structure and fix helical defects (Tab. S4). For each backmapped replicate 2 *µs* of simulation were conducted (12 × 2 *µs*).

### Enhanced sampling simulations

Enhanced sampling simulations were run with the initial all-atom box of the Bcl-xL/tBid complex. To accelerate the orientation of Bcl-xL’s globular domain on top of the membrane the torsion between the globular domain and the z-axis along the tBid BH3 helix was biased with OPES explore [42]. Additional walls were applied to the system to retain protein fold, orientation of the tBid BH3 helix, and disallow membrane deformation. Three independent replicates were simulated for 1 *µs*. From them the free energy landscape along the torsion angle was estimated via reweighting and kernel density estimation. In estimation the initial 200 *ns* were discarded and bandwidth selected with the Silverman’s rule of thumb. The free energy landscape was also calculated for the re-seeded all-atom simulations to validate position of the identified global minima of globular domain orientation (Fig. S13E).

### All-atom molecular dynamics analysis

Production simulations (both initial, Fig. S11, and re-seeded) were analysed visually with VMD and pymol [64, 84]. Root mean squared deviation (RMSD), root mean squared fluctuation (RMSF), atom based principle component analysis (PCA), solvent accessible surface area (SASA), angles and distances were calculated using GROMACS tools and MDanalysis [85]. Unless specified otherwise, trajectories were analysed at an interval of 1 *ns*^−1^. Both for RMSF and atom based PCA analysis simulations were aligned to the first frame of the simulation based on the C*α*-atoms of helices *α*5 and *α*6 and analysed at 10 *ns*^−1^. The PCA aligned trajectory was also used to compute the dynamic cross-correlation matrix of C*α*-atom displacement of protein residues.

DEER distributions within re-seeded all-atom simulations were calculated using DEER-PREdict [41] (see: equation 0.2) for each frame in the trajectories (10 *ns*^−1^). Total distributions were obtained by summing the distributions of separate trajectories and normalising by *∫ P* (*d*).

HDX deuteration levels in re-seeded all-atom simulations were calculated using HDXer^2^ [86, 87]. Simulations were analysed with the method of Best and Vendruscolo [87] at deuteration times of 0.05*/*0.5*/*5.0 *min*, 22^◦^*C*, at pH 7.5 and maximum deuteration fraction of 90.4 %.

Contact maps and probabilities were calculated using custom wrapper modules around MDAnalysis [85] defining a contact as a distance lower than 0.7 *nm* between the relevant heavy side-chain atom center of mass (CoM) of two protein residues. Independently moving domains within the hetero-complex were identified via the standard deviation matrix of distances between the centres of mass of all protein residue side-chains. Membrane contacts were calculated at 0.5*ns*^−1^ between protein residues and membrane phosphates at a cutoff of 1 *nm*. Binding site dynamics were compared by calculating the distance between the centre of mass of the backbone of residues 107 to 112 and 137 to 141 in Bcl-xL. The angle of the binding site in relation to the membrane was calculated as the angle between the z-axis and the vector between the *Cα* atoms of A104 and E129.

To obtain an overview over loop and general dynamics of the complex in the two sets of all-atom simulations (initial, re-seeded) a PCA was calculated for the distance matrix between Bcl-xL and tBid using the same selections as for the contact maps (Fig. S14). The PCA explained 90 % of cumulative variance in the distance matrix between Bcl-xL and tBid and was consecutively clustered first by OPTICS and then by KMeans to obtain the number of clusters and assign a state to all considered frames. Clusters were sorted by size.

All structural images were rendered with PyMOL [64] or Molecular Nodes^3^ in Blender. Biological graphics were supplemented with visual’s obtained from BioRender^4^.

## Supporting information

SI material for MD simulations and experiments as referenced in the text

## Acknowledgements

Technical support for protein purifications was provided by R. Visentin, L. Falconet (Protein Platform, Faculty of Medicine, University of Geneva) and A. Cieren (Department of Physical Chemistry, University of Geneva). The mouse livers were kindly provided by Service de Zootechnie (Faculty of Medicine, University of Geneva) and the mitochondria were prepared by L.M. Epasto and A. Cieren (Department of Physical Chemistry, University of Geneva). Technical support for HDX-MS was provided by R. Visentin (Protein Platform, Faculty of Medicine, University of Geneva). EB and CE thank Prof. A.J. García-Sáez for providing the Bcl-2 plasmids and Prof. F. Marassi for providing the pdb files of full-length soluble and membrane-anchored Bcl-xL. AH and FLG thank Dr. N. Piasentin and Dr. S. Aureli for reading and commenting on the manuscript. This work is funded by the SNSF Sinergia grant CRSII5_216587 (to E.B. and F.L.G).

## Author Contributions and Declarations

EB and FLG conceived the project with the support of CE and AH. CE, AH, OV conducted experiments and analysis. CE conducted DEER, ESEEM and protein preparations. AH conducted modelling and simulation. OV conducted HDX experiments. CE, AH, FLG and EB conceptualised and wrote the manuscript. EB and FLG acquired funding.

This work was supported by the SNSF Sinergia grant CRSII5_216587 (to EB and FLG) and partially by grant 200021_204795 (to FLG). We would like to acknowledge the CSCS for their generous allocation of supercomputing time (project: lp84). The authors declare no competing interests.

## Data Availability

MD simulations, modeller and analysis code will be accessible on Zenodo under following DOI: https://doi.org/10.5281/zenodo.16941554 upon publication. Experimental data such as ESEEM, DEER and HDX will be accessible on Zenodo under following DOI: https://doi.org/10.5281/zenodo.16941554 upon publication.

## Footnotes

1 such that *∫ P (d)* = 1.0

2 https://github.com/Lucy-Forrest-Lab/HDXer

3 https://github.com/BradyAJohnston/MolecularNodes

4 https://www.biorender.com/

