## Supplementary material for "Structural and dynamic basis of indirect apoptosis inhibition by Bcl-xL: a case study with Bid": SI material for MD simulations and experiments as referenced in the text

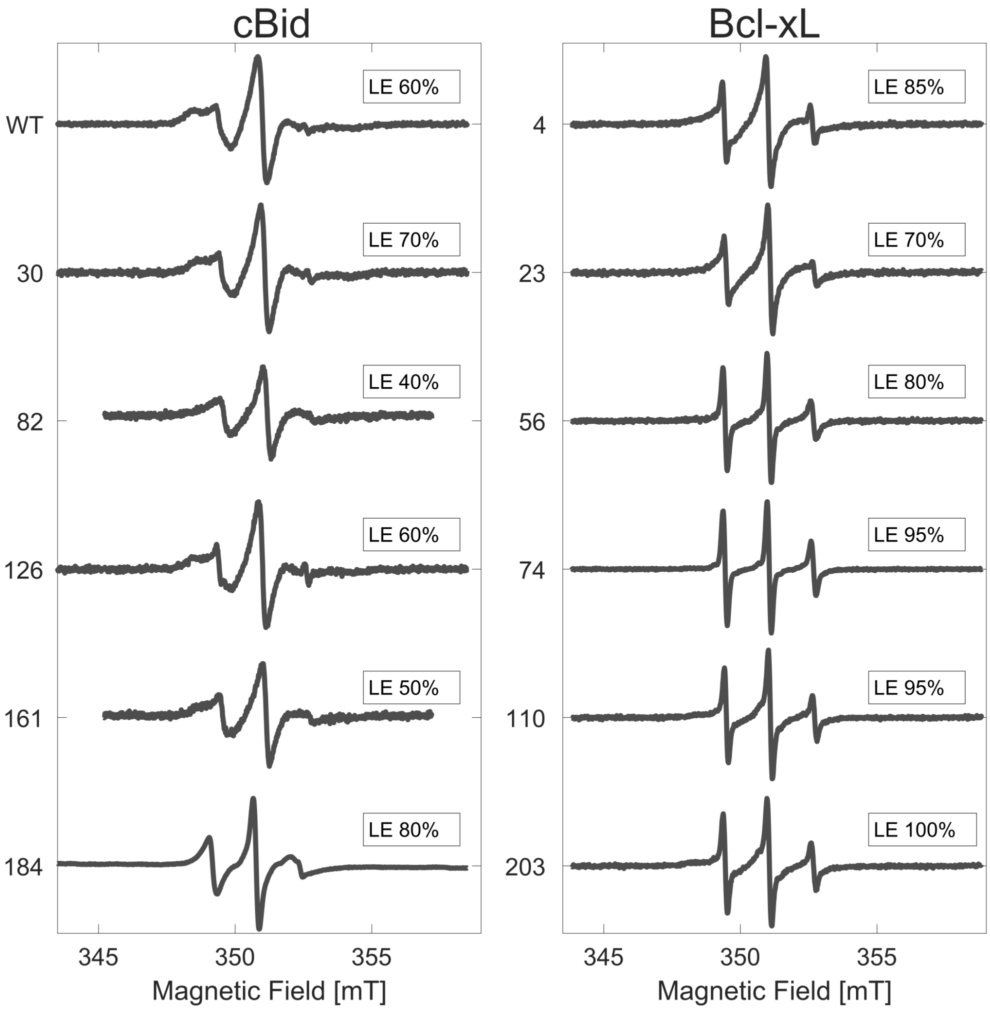


**Figure 1: Room temperature continuous wave EPR spectra of spin-labeled cBid and Bcl-xL variants.** The wild type cBid (WT) contains two natural cysteines (C30 and C126) which were both spin labeled. The other cBid variants are singly spin-labeled at the indicated cysteine positions (one or two natural cysteines were replaced by serines). The Bcl-xL variants are singly spin-labeled at the indicated cysteine positions (the natural cysteine C151 was replaced by serine). Labeling efficiencies (LE, spin per cysteine, 10% error) were determined by comparing the second integral of the spectra with the spectrum of TEMPOL in water with known concentration.

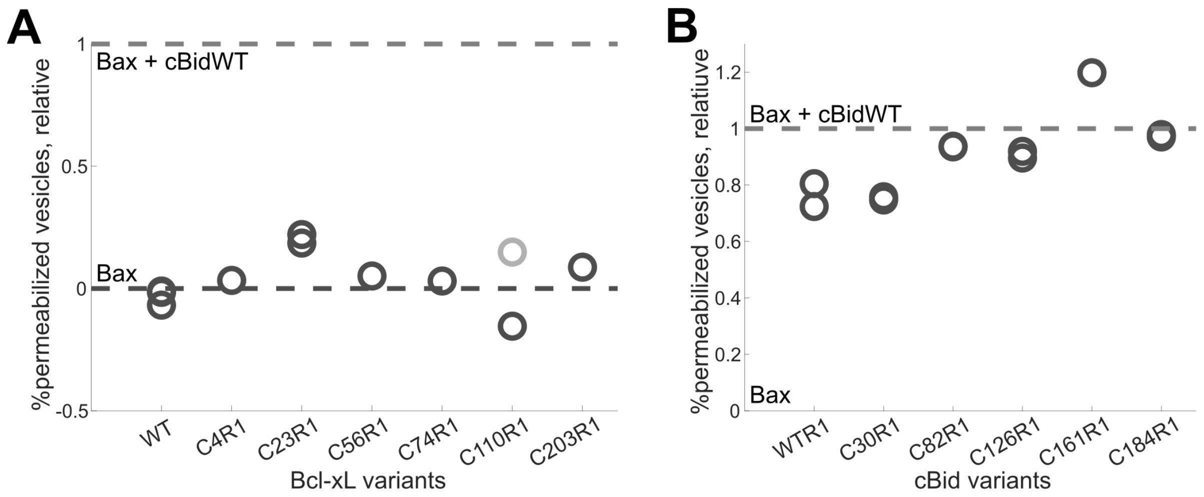


**Figure 2: Activity of spin-labeled Bcl-xL and cBid variants by pore-formation assays.** Comparison of the release of calcein from LUVs for different Bcl-xL **(A)** and cBid **(B)** spin-labeled variants after 30 min incubation. **(A)** Activity of Bcl-xL variants in presence of Bax and cBid (25 nM concentrations 1:1:1 protein ratio). To facilitate the comparison of the inhibitory effects of Bcl-xL variants with different LUVs batches, data were normalized as follows: zero is the calcein release with Bax alone (Bax autoactivity) and one the maximum release obtained with Bax and cBidWT. All spin-labeled variants are shown to inhibit membrane permeabilization. **(B)** Activity of cBid variants in presence of Bax (25 nM concentrations 1:1 protein ratio). Normalization as in panel A.

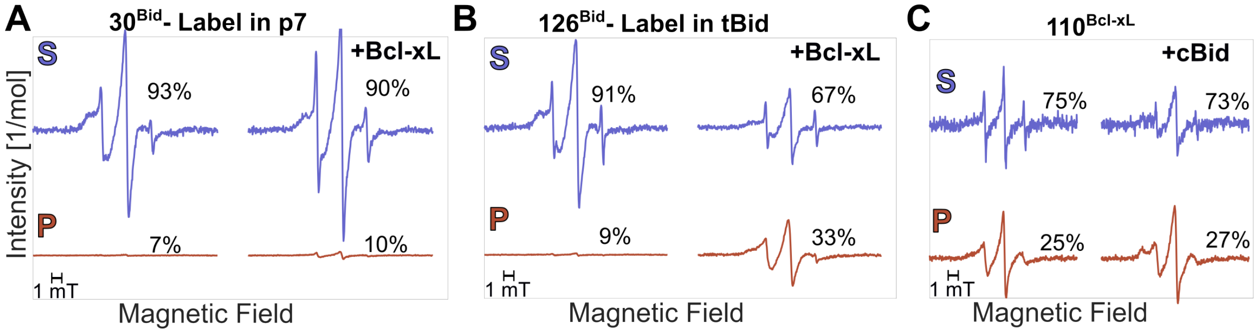


**Figure 3: Membrane-water partitioning of spin-labeled cBid and Bcl-xL variants.** Room temperature continuous wave EPR spectra of spin-labeled cBid and Bcl-xL variants were recorded in the supernatant (S) and pellet (P) fractions after incubation. The number of moles of p7 **(A)**, tBid **(B)** and Bcl-xL **(C)** in each fraction was calculated based on the spectral area (spin concentration) and the volume of each fraction. The spectra are normalized by the number of moles in each fraction and the relative amount is shown in percentage (the estimated error is 10%), presented in fig. 2B

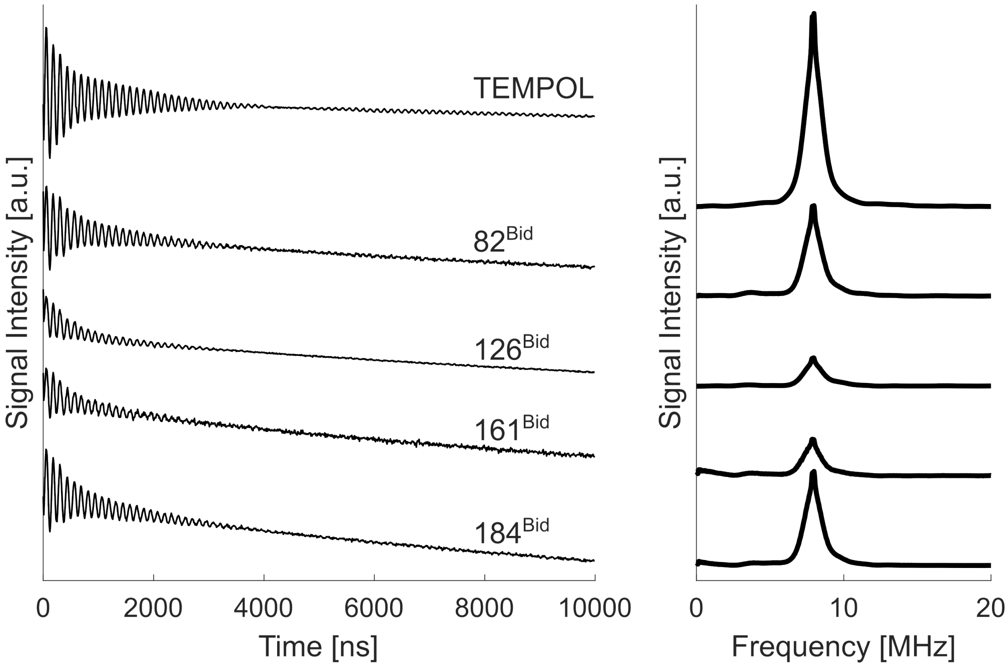


**Figure 4: 3-pulse ESEEM data of spin-labeled cBid in presence of unlabeled Bcl-xL in pellet fractions.** Pellets were resuspended with deuterated buffer and 10% $d_8$-glycerol. Left: Raw data of Fig. 2C normalized by the number of scans. Right: Deuterium signal intensity after Fourier transformation, presented in fig. 2C (the estimated error based on technical repeats is 10%). The reference sample was prepared by adding TEMPOL after pellet resuspension.

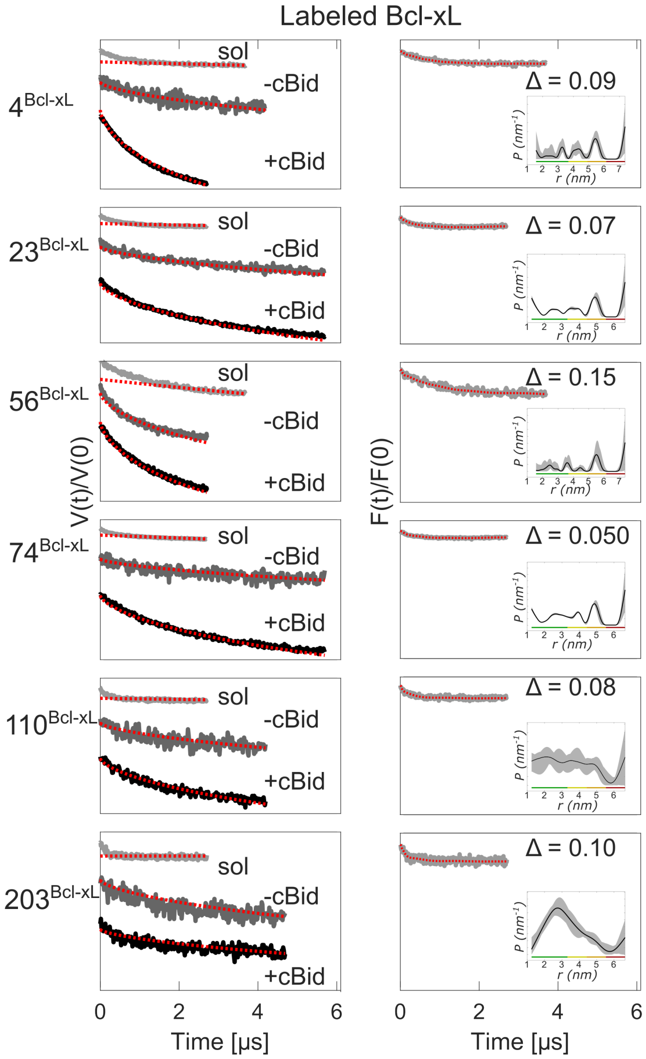


**Figure 5: DEER analysis of singly-labeled Bcl-xL variants in solution (sol) and in LUVs without cBid wt (-cBid), and in LUVs with unlabeled cBid wt (+cBid).** Left panels: Primary data (grey and black) and 3D background fit (dotted red, for the samples in solution - sol-) or 2D background fit (dotted red, for the membrane samples -cBid and +cBid). The 2D-background (starting at the zero time point) is almost indistinguishable from the DEER primary traces in membranes, indicating that Bcl-xL does not form defined dimers or higher order oligomers. The background is often steeper when adding cBid which might indicate higher molecular crowding when tBid is bound to Bcl-xL at the membrane. Right panels: Form factor of the solution samples obtained after dividing the primary data by the background (grey), fit (red dotted) and corresponding modulation depth (Δ). The insets show the resulting distance distributions obtained with Tikhonov regularization. the low modulation depth and the broad distributions are characteristic of residual Bcl-xL/Bcl-xL aggregates in solution.

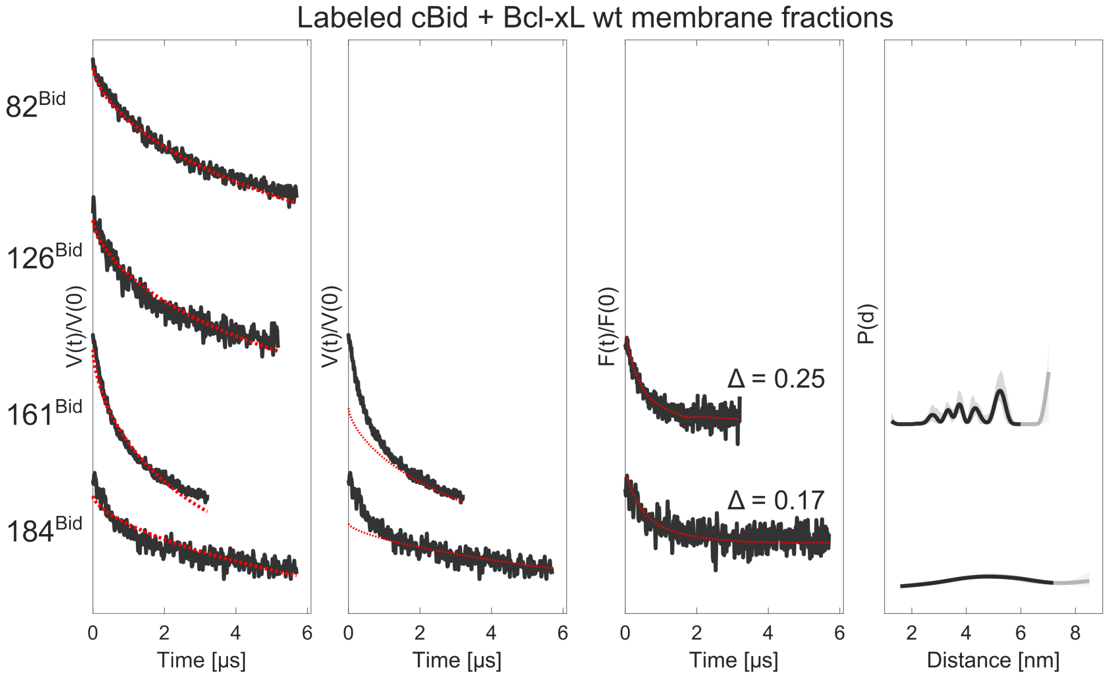


**Figure 6: DEER analysis of singly-labeled cBid variants in LUVs unlabeled Bcl-xL wt.** From left to right: Primary data (black) with a 2D background fit (dotted red) starting from zero; the same primary data of two traces with a 2D background starting at 2 microseconds (dotted red); Form factor obtained after dividing the primary data by the background (black), fit (red dotted) and corresponding modulation depth (Δ); resulting distance distributions obtained with Tikhonov regularization with uncertainties (shaded grey areas) derived by varying the starting point of the background fit and the noise level. The comparison of the traces with a decay caused only by the two dimensional distribution of the spin-labeled proteins highlights that the traces are almost indistinguishable from a 2D background, indicating that there are no specific cBid/cBid interactions at the membrane when the inhibitory complex is formed. For positions 161 and 184 the traces deviate more from from a 2D background. The fit with a 2D background starting at 2 microsecond demonstrate that aspecific interaction are detected, indicating molecular crowding at the membrane of the C-terminal region of cBid.

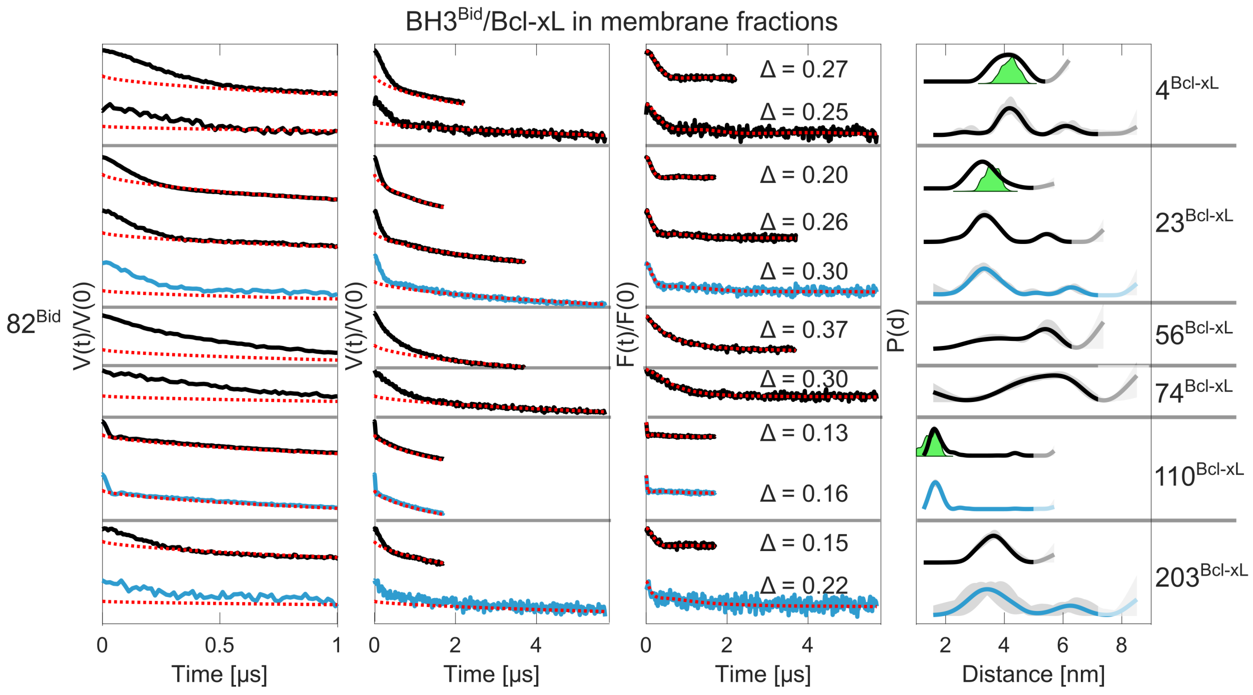


**Figure 7: BH3$^{Bid}$/Bcl-xL distance measurements in membranes (part I).** From left to right: One microsecond zoom on the primary DEER data (black) with a 2D background fit (dotted red); primary DEER data (black) with a 2D background fit (dotted red); the same primary data of two traces with a 2D background starting at 2 microseconds (dotted red); Form factor obtained after dividing the primary data by the background (black), fit (red dotted) and corresponding modulation depth (Δ); resulting distance distributions obtained with Tikhonov regularization with uncertainties (shaded grey areas) derived by varying the starting point of the background fit and the noise level. Technical and biological repeats are shown. The traces are shaded to highlight the distance range in which the accuracy is not adequate, due to the maximal length of the trace detected. The blue traces are those used as first three constraints to create the initial model in Fig. 3. The three green areas represent the three distance distributions which can be predicted with MMM[1] on the crystal structure of the water-soluble truncated Bcl-xL in complex with a Bid-derived BH3 peptide (PDB: 4QVE). The predicted distributions are in very good agreement with the distributions obtained with the full-length Bcl-xL and cBid at the membrane, confirming the presence of the same BH3-into-groove interaction.

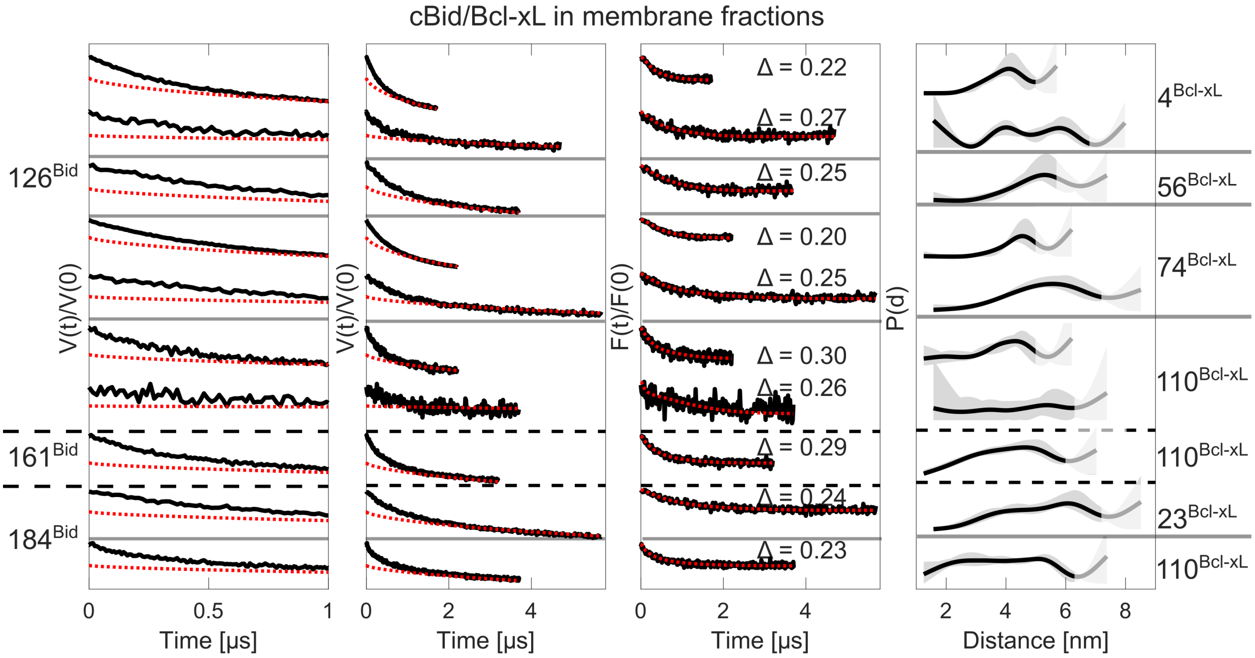


**Figure 8: cBid/Bcl-xL distance measurements in membranes (part II).** From left to right: One microsecond zoom on the primary DEER data (black) with a 2D background fit (dotted red) ; primary DEER data (black) with a 2D background fit (dotted red); the same primary data of two traces with a 2D background starting at 2 microseconds (dotted red); Form factor obtained after dividing the primary data by the background (black), fit (red dotted) and corresponding modulation depth (Δ); resulting distance distributions obtained with Tikhonov regularization with uncertainties (shaded grey areas) derived by varying the starting point of the background fit and the noise level. The traces are shaded to highlight the distance range in which the accuracy is not adequate, due to the maximal length of the trace detected. For the same sample, traces of different lengths are shown as technical repeats.

```
E-value: 1e-73  Identity: 126/198
Cleave site indicated by: xxxx (cleaved after last x)
                                                                  xx
Human    MDCEVNNGSSLRDECITNLLVFGFLQSCSDNSFRRELDALGHELPVLAPQWEG--YDELQ
Mouse    MDSEVSNGSGLGAEHITDLLVFGFLQSSG--CTRQELEVLGRELPVQA-YWEADLEDELQ
         **.**.***.*  * **:*********..  . *:**:.**:**** *  **.   ****
         xx
Human    TDGNRSSHSR-LGRIEADSESQEDIIRNIARHLAQVGDSMDRSIPPGLVNGLALQLRNTS
Mouse    TDGSQASRSFNQGRIEPDSESQEEIIHNIARHLAQIGDEMDHNIQPTLVRQLAAQFMNGS
         ***.::*:*   ****.******:**:********:**.**:.* * **. ** *: * *

Human    RSEEDRNRDLATALEQLLQAYPRDMEKEKTMLVLALLLAKKVASHTPSLLRDVFHTTVNF
Mouse    LSEEDKRNCLAKALDEVKTAFPRDMENDKAMLIMTMLLAKKVASHAPSLLRDVFHTTVNF
          ****:.. **.**:::  *:*****::*:**::::*********:**************

Human    INQNLRTYVRSLARNGMD    " ": different
Mouse    INQNLFSYVRNLVRNEMD    ".": similar residue
         ***** :***.*.** **    ":": highly similar residue
```

**Figure 9: Sequence alignment of human and mouse Bid.** Human tBid was used for the modeling, mouse cBid was used for the EPR and HDX-MS experimental data. The cleavage occurs at the end of the sequence LQTD (indicated with xxxx), with D being at position 59 for mouse and 60 for human Bid. The N-terminal fragment of cBid is called p7 (7 kDa) and the C-terminal fragment tBid (15 kDa). Alignment generated with clustalw.

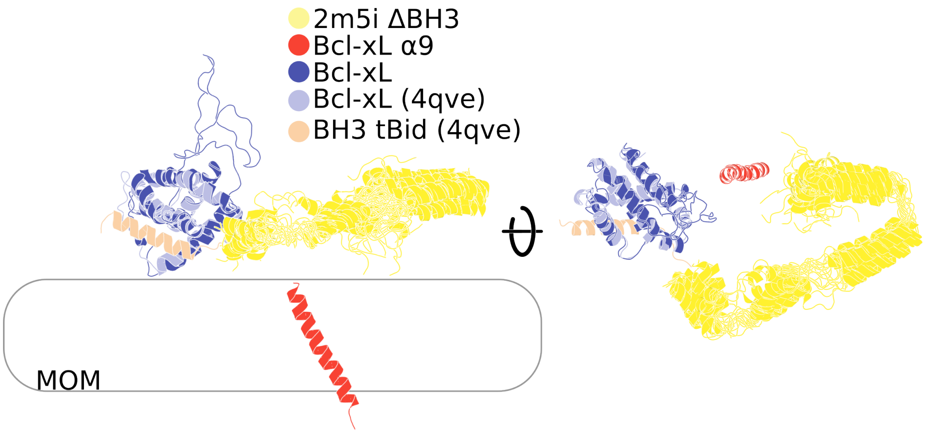


**Figure 10: Positioning of input structures for Modeller shown as cartoon.** The Bcl-xL input structure was obtained from [2] and positioned through alignment to the globular domain of PDB 4qve [3]. For details on sub-structure placement and generation see Model generation in Methods. Sub-structures of the aligned input structures were combined into the starting models with Modeller [4] in a single step (see Model generation in Methods).

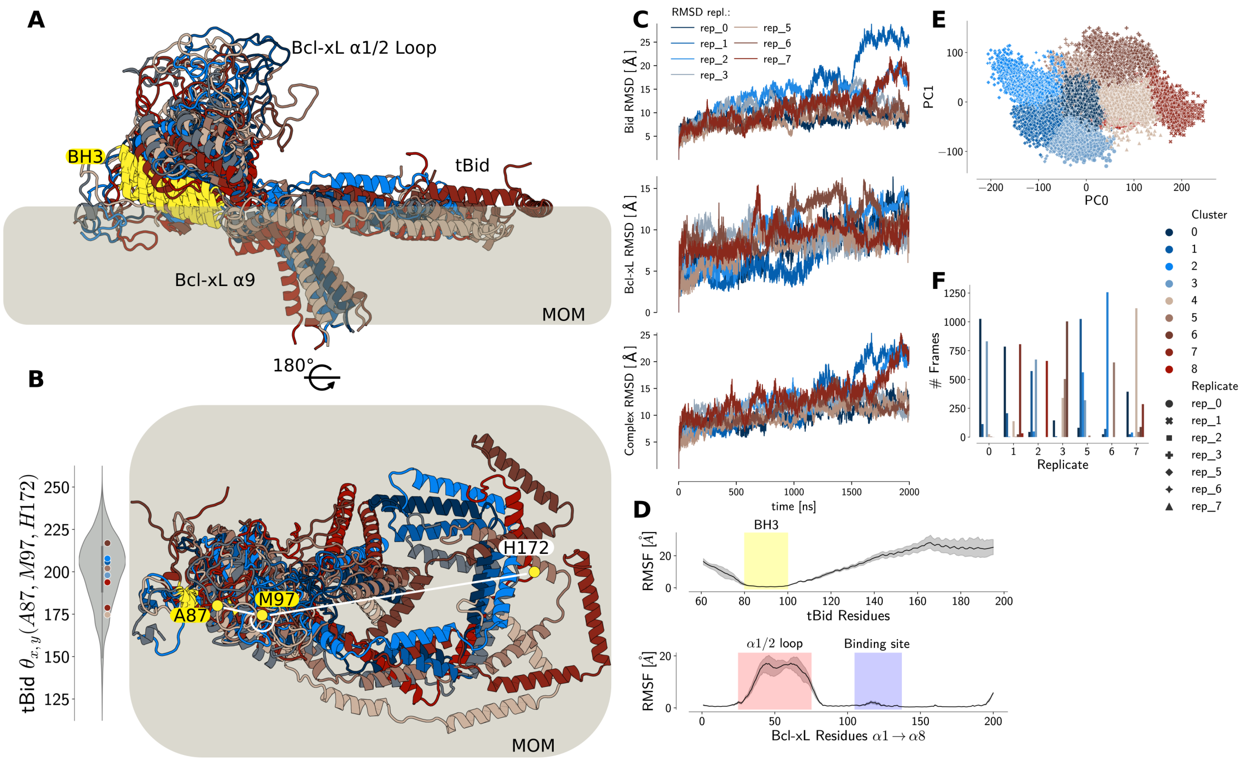


**Figure 11: Analysis of initial $7 \times 2\,\mu s$ of all-atom MD. (A)** Cluster representatives of the Bcl-xL/tBid complex structural ensemble embedded on a lipid membrane (cartoon, side view). The tBid BH3 domain is shown in yellow. Cluster representatives were used to seed coarse grained Martini simulations. **(B)** tBid's helices $\alpha 4-8$ diffuse freely on the membrane (cartoon, top view), as shown by their angular spread with respect to the BH3 domain (angle-defining atoms as yellow spheres). **(C)** RMSF of tBid and Bcl-xL. Structures were aligned on the $\alpha 5$ and $\alpha 6$ helical C$\alpha$-atoms of Bcl-xL. Specific regions of the sequences are highlighted. **(D)** RMSD of independent simulation replicates over time, decomposed into the complex and its components. **(E)** Clustering of frames based on distances between Bcl-xL and tBid mapped onto the first two principal components (see methods). **(F)** Decomposition of clusters and their sizes over the different independent simulation replicates.

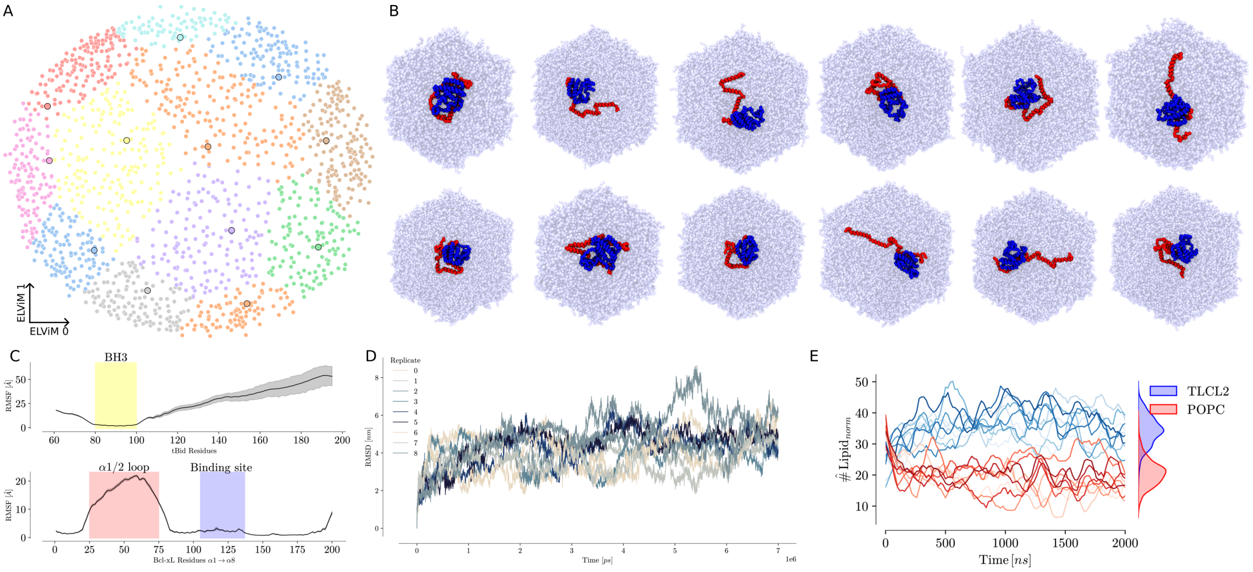


**Figure 12: Coarse grained simulations.** **(A)** ELViM projection of the CG-MD ensemble based on the backbone beads and clustered via Gaussian mixture models (12 clusters). Cluster representatives indicated as larger dots with black outlines. **(B)** Cluster representatives extracted from the Gaussian mixture model used to re-seed all-atom MD simulations. Membrane shown as transparent blue beads, Bcl-xL as blue beads and tBid as red beads. **(C)** Average RMSF of the CG-MD simulations for tBid (top) and Bcl-xL(bottom). **(D)** RMSD of the independent CG-MD simulations over time, colored by replicate. **(E)** Cardiolipin enriches around the complex within CG-MD simulations across all replicates.

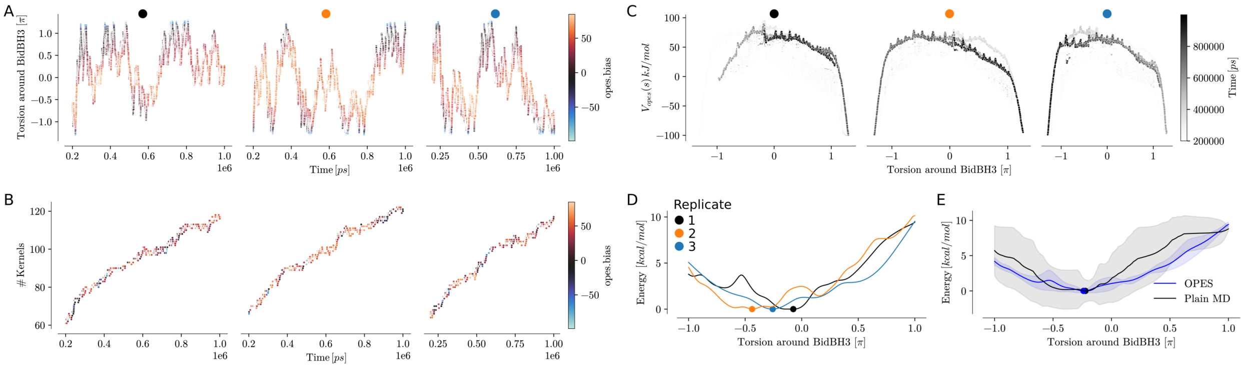


**Figure 13: Opes explore simulations.** **A** Sampling in the independent replicates of the accelerated torsion over time. This torsion constitutes the orientation of the Bcl-xL binding domain relative to the membrane normal over tBid's BH3 helix. **B** Number of kernels used to compute the bias potential ($V_t(s)$) at time $t$. **C** Shape of the bias potential $V_t(s)$ over the accelerated torsion colored over time. **D** Free energy surfaces (FES) computed for the independent replicates. Global minima indicated as dots. **E** Average Opes explore FES compared to the average FES of the re-seeded all-atom MD simulations. Global minima indicated as dots.

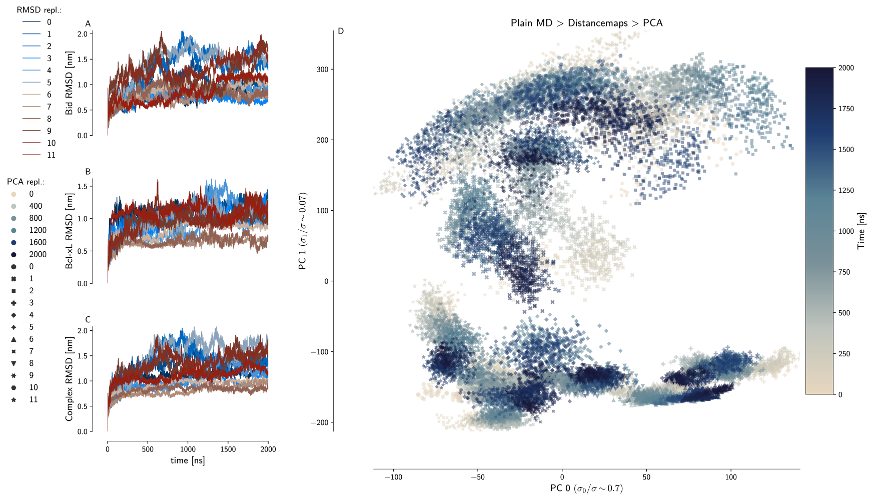

**Figure 14: RMSD and PCA of re-seeded all-atom MD simulations.** (A-C) RMSD of independent simulation replicates over time, decomposed into the Complex (C) and its components (A, B). (D) Scatter plot of the independent simulation replicates within the two first principle components of distances between Bcl-xL and tBid $C\alpha$-atoms.

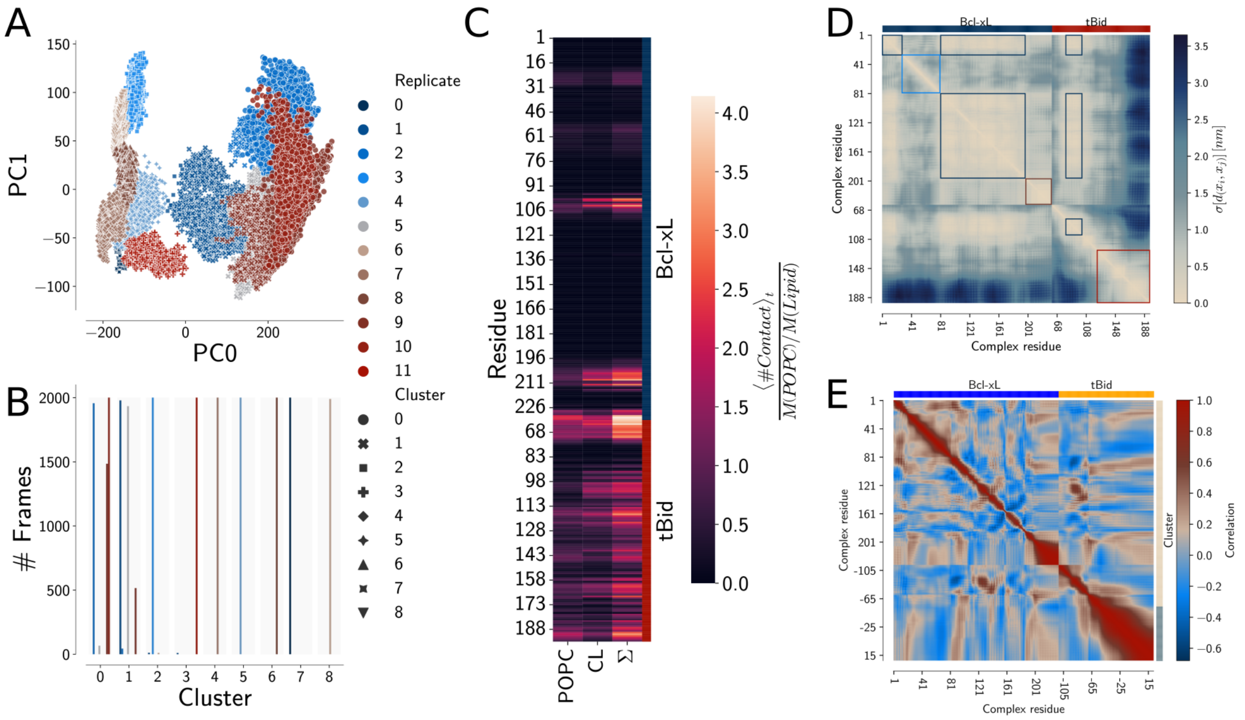


**Figure 15: Supplementary materials for Fig. 4.** **(A)** Clustering of frames based on distances between Bcl-xL and tBid mapped onto the first two principal components (see methods). **(B)** Decomposition of clusters and their sizes over the different independent simulation replicates. **(C)** Contact frequency of each residue in Bcl-xL and tBid with either POPC or Cardiolipin (CL). Contacts were calculated between protein residue and lipid phosphates center of masses with a radius of $1\,nm$. Frequencies were mass normalized, to consider the bulkiness of cardiolipin compared to POPC. **(D)** Standard deviation matrix of protein residue residue distances, the identification of the different clusters (squares with the same color) is guided by biological intuition. **(E)** Dynamic cross-correlation matrix of residue displacement (blue to red) clustered by agglomerative clustering (right heatmap margin; gray and beige) with the cluster number chosen based on the highest silhouette score ($\sim 0.32$). This clustering identifies separately the membrane diffusing C-terminal helices of tBid and water diffusing parts of the complex, composed of the globular Bcl-xL domain, $\alpha 1/2$ loop and tBids BH3 helix. The clustering groups Bcl-xL's TMD with the water diffusing cluster, due to its proximity to the globular domain and diffusion limited by the C-terminal helices of tBid when these wrap around the complex.

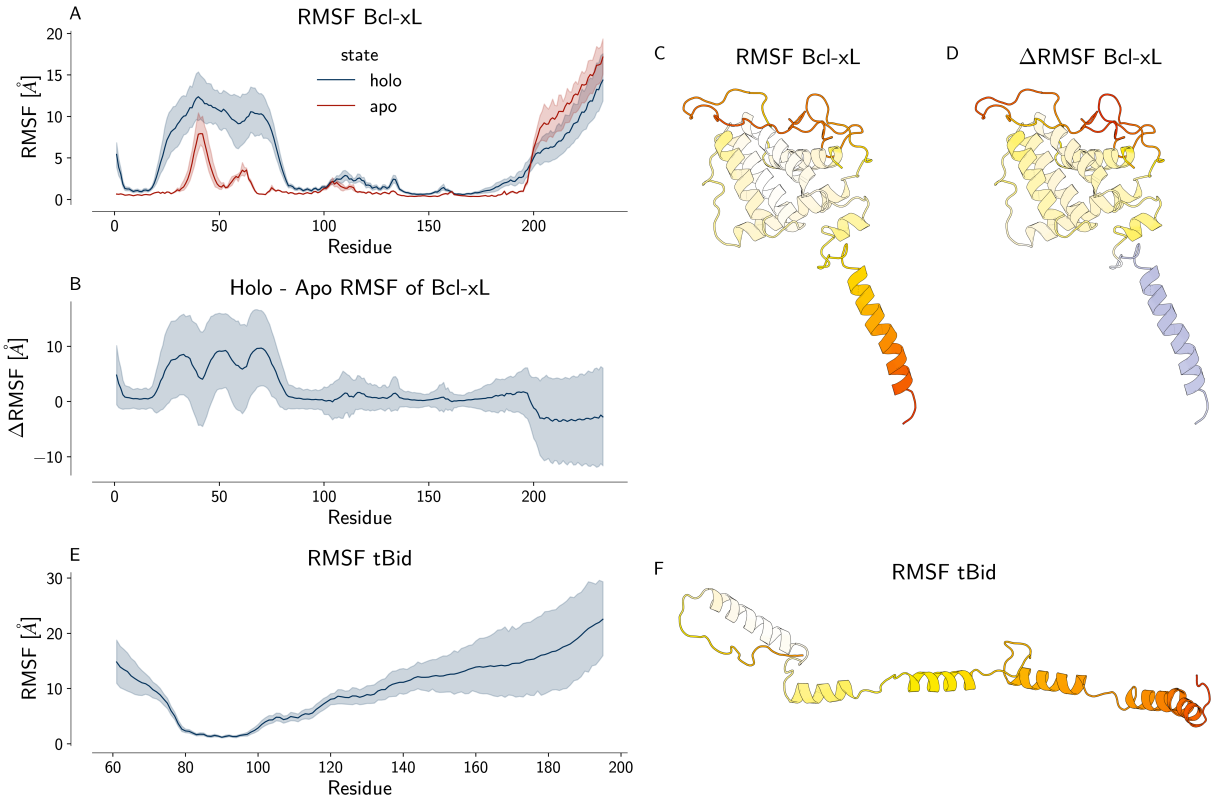


**Figure 16: Binding of tBid to Bcl-xL increases Bcl-xL's flexibility.** **(A)** RMSF comparison of the apo and holo simulations of Bcl-xL. **(B)** ΔRMSF of apo and holo Bcl-xL. **(C)** Cartoon representation of Bcl-xL with the holo RMSF mapped (low: white; high:red) onto the structure, note the highest RMSF is in the $\alpha 1/2$ loop and $\alpha 9$ (trans membrane domain). **(D)** Cartoon representation of Bcl-xL with the ΔRMSF mapped (negative: blue; low: white; positive: red) onto the structure. The highest impact of binding onto Bcl-xL flexibility is in the $\alpha 1/2$ loop. **(E)** RMSF of tBid in the hetero-complex. **(F)** Cartoon representation of tBid on the membrane colored by RMSF (low: white; high: red), note that the BH3 domain of tBid is rigidly bound to the globular domain of Bcl-xL. RMSF values were calculated by aligning all simulations to the residues 54 to 195 within Bcl-xL in their first frame.

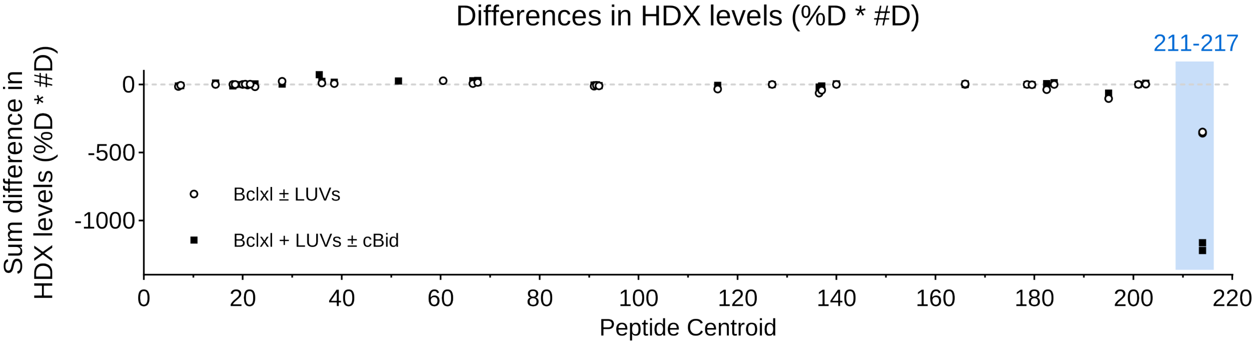


**Figure 17: Differences in deuteration levels of Bcl-xL: Comparison of deuterium incorporation for each studied peptide is represented.** The sum of differences for all timepoints is calculated, both as percentage deuteration ($\%D$) and number of deuterons ($\#D$). The product of the sum in differences in $\%D$ and $\#D$ is then multiplied to increase signal-to-noise ratio. Regions highlighted in blue indicate protection from exchange by the ligand. White circles: Bcl-xL in LUVs compared to Bcl-xL in solution. Black squares: Bcl-xL in LUVs with cBid compared to Bcl-xL alone in LUVs.

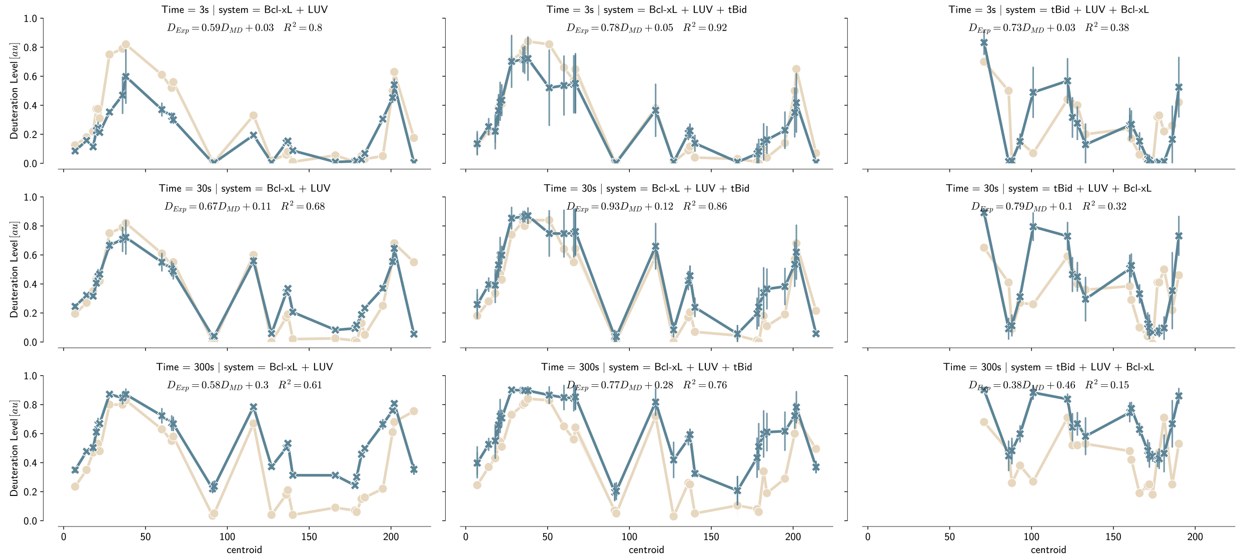


**Figure 18: Deuteration levels for both Bcl-xL and Bcl-xL/tBid on LUV's.** Comparison between experimental measurements (Bcl-xL + mouse cBid) and MD simulations (Bcl-xL + human tBid). Values calculated as mean across re-seeded all-atom simulations with standard deviations indicated as error bars. $R^2$ and linear equations are obtained by linear regression of the experimental against the average re-seeded all atom MD deuteration factors.

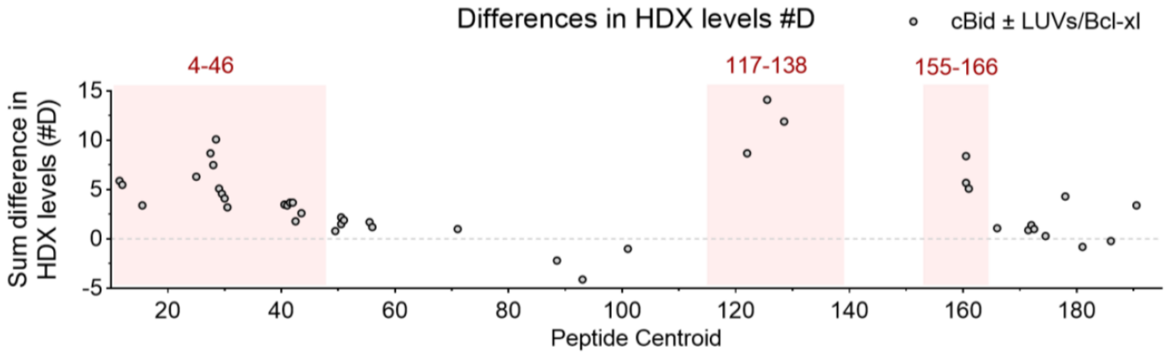


**Figure 19: Comparison of the deuterium incorporation for each studied peptide of cBid in the pellet fraction (cBid mixed with Bcl-xL and LUVs) versus cBid alone in solution.** The sum of differences (pellet minus solution) for all time points is calculated, both as percentage deuteration ($\%D$) and number of deuterons ($\#D$). The product of the sum in differences in $\%D$ and $\#D$ is then multiplied to increase signal-to-noise ratio. Regions highlighted in red have a higher deuterium exchange going from the soluble cBid to the membrane-associated complex. cBid is cleaved by caspase 8 after Asp 60 in mouse Bid.

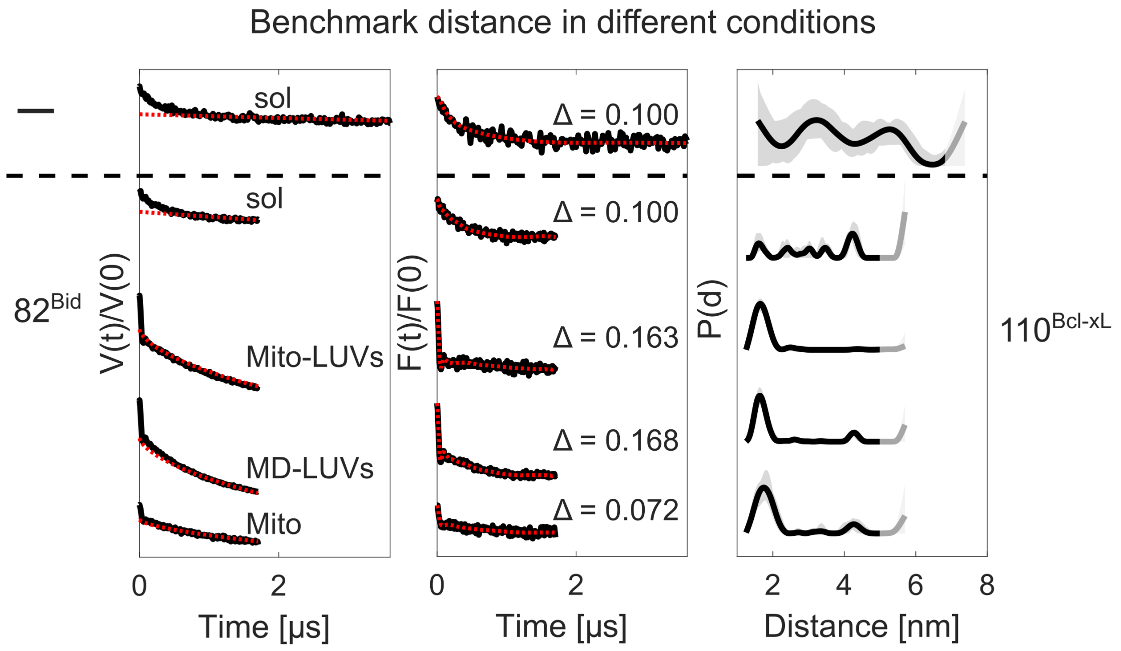


**Figure 20: Benchmark $82^{\mathbf{Bid}}/110^{\mathbf{Bcl\text{-}xL}}$ distance measurements in different conditions.** From left to right: Primary DEER data (black) with 3D- (for solution samples) or 2D-background fit (red dotted); Form factor obtained after dividing the primary data by the background (black), fit (red dotted) and corresponding modulation depth (Δ); Resulting distance distributions obtained with Tikhonov regularization with uncertainties (shaded grey areas) derived by varying the starting point of the background fit and the noise level. The traces are shaded to highlight the distance range in which the accuracy is not adequate, due to the maximal length of the trace detected.

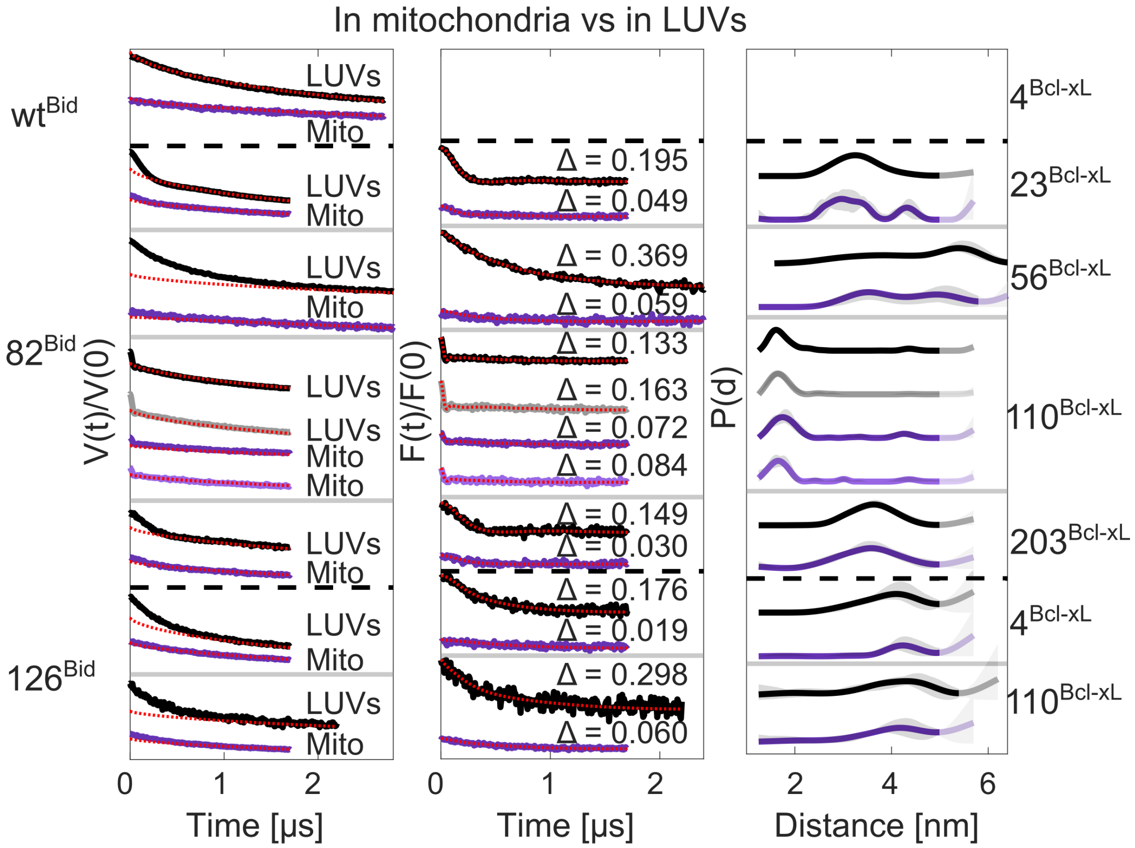


**Figure 21: Comparison of *in vitro* and *in organelle* distance measurements.** From left to right: Primary DEER data (black LUVs, purple mitochondria) with 2D-background fit (red dotted); Form factor obtained after dividing the primary data by the background (black/purple), fit (red dotted) and corresponding modulation depth (Δ); Resulting distance distributions obtained with Tikhonov regularization with uncertainties (shaded grey areas) derived by varying the starting point of the background fit and the noise level. The traces are shaded to highlight the distance range in which the accuracy is not adequate, due to the maximal length of the trace detected. Biological repeats are shown.

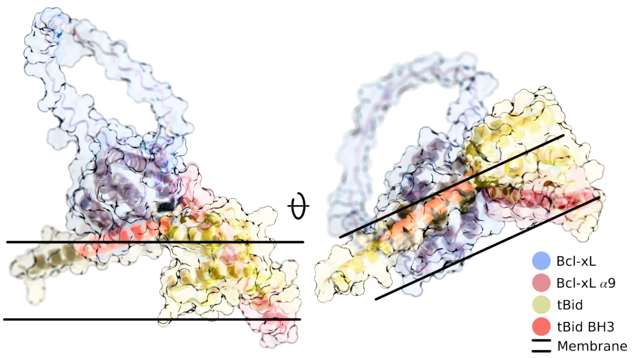


**Figure 22: Alphafold model of the Bcl-xL/tBid complex.** Protein complex shown as cartoon and surface colored by domain (globular Bcl-xL domain, TMD, tBid BH3 and tBid). The two parallel lines guide the eyes to indicate a putative membrane bilayer in which the transmembrane helix of Bcl-xLis inserted. Note the folding of tBid around the TMD of Bcl-xL, which would impede a correct membrane insertion of the transmembrane helix of Bcl-xLin this model. Models were generated with ColabFold2.

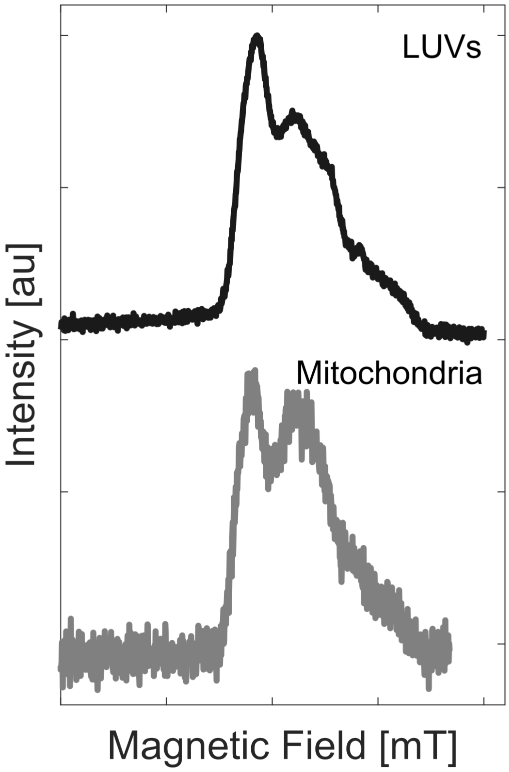


**Figure 23: Field swept echo of LUV vs mitochondria samples showing the effects of metals on the field sweep.** Sample: $110^{\text{Bcl-xL}} + 82^{\text{Bid}}$ pellet samples with either LUVs or mitochondria. Shot repetition time: 1 ms, $t_1$: 1000 ns, $t_2$: 400 ns, Sweep width: 20 mT, $\frac{\pi}{2}$-pulse: 18 ns, $\pi$-pulse: 36 ns.

**Table 1: Tabulated contacts between tBid and Bcl-xL in MD simulations.** Contact frequency was calculated as the ratio of MD frames ($ns^{-1}$) where the contact was observed. Contacts are defined as a distance lower than 7 Å between the CoM of residue sidechains including hydrogen's. Relevant non-hydrophobic interactions are indicated with red dots.

| | tBid | | Bcl-xL | | |
|---|---|---|---|---|---|
| | Residue name | Residue position | Residue name | Residue position | Contact frequency |
| | Constant contacts $f_c > 0.8$ | | | | |
| | ILE | 83 | SER | 122 | 0.872 |
| | ILE | 83 | VAL | 126 | 0.893 |
| | ALA | 87 | VAL | 126 | 0.977 |
| | LEU | 90 | ALA | 142 | 0.937 |
| | LEU | 90 | PHE | 146 | 0.972 |
| | ALA | 91 | LEU | 130 | 0.876 |
| | ALA | 91 | ARG | 139 | 0.867 |
| | VAL | 93 | PHE | 97 | 0.949 |
| | GLY | 94 | ALA | 142 | 0.959 |
| • | ASP | 95 | ARG | 139 | 0.965 |
| | MET | 97 | PHE | 97 | 0.893 |
| | Transient contacts $0.5 < f_c < 0.8$ | | | | |
| | ILE | 83 | GLN | 125 | 0.606 |
| • | ARG | 84 | GLU | 129 | 0.725 |
| | ILE | 86 | LEU | 108 | 0.539 |
| | ILE | 86 | LEU | 112 | 0.695 |
| | ILE | 86 | VAL | 126 | 0.562 |
| | ILE | 86 | PHE | 146 | 0.793 |
| | ALA | 87 | GLU | 129 | 0.755 |
| | ALA | 87 | LEU | 130 | 0.796 |
| | HIS | 89 | PHE | 105 | 0.595 |
| | LEU | 90 | VAL | 126 | 0.607 |
| | LEU | 90 | LEU | 130 | 0.795 |
| | VAL | 93 | TYR | 101 | 0.798 |
| | VAL | 93 | PHE | 105 | 0.503 |
| | VAL | 93 | ALA | 142 | 0.781 |
| | GLY | 94 | PHE | 97 | 0.689 |
| | GLY | 94 | GLY | 138 | 0.798 |
| • | ASP | 95 | ASN | 136 | 0.549 |
| | MET | 97 | ALA | 93 | 0.616 |
| | MET | 97 | GLY | 138 | 0.785 |
| | MET | 97 | VAL | 141 | 0.706 |
| • | ASP | 98 | ASN | 136 | 0.561 |
| | ASP | 98 | GLY | 138 | 0.696 |

**Table 2: Applied DEER parameters.**

| | NO-NO |
|---|---|
| Shot repetition time [ms] | 5 |
| $t_1$ [ns] | 400 |
| $t_2$ [ns] | 2000 - 6000 |
| Zero time [ns] | 120 |
| $t_1$ averages in steps of [ns] | 8 or 16 |

**Table 3: Applied ESEEM parameters.**

| | |
|---|---|
| Video band width [MHz] | 20 |
| Shot repetition time [ms] | 5 |
| $\tau$ [ns] | 304 |
| T start [ns] | 20 |
| T increment [ns] | 10 |
| T increment steps | 1000 |

**Table 4: Simulation protocol and restraint used throughout.** For detailed description on other simulation parameters see Methods.

| System | integrator | nsteps | emtol | constraints | constraint algorithm | replicates |
|---|---|---|---|---|---|---|
| initial Holo | steep | 50000 | 1000.0 | – | – | 1 |
| martini Holo | steep | 5000 | 1000.0 | – | – | 9 |
| re-seeded Holo | steep | 50000 | 1000.0 | – | – | 12 |
| Apo | steep | 5000 | 1000.0 | h-bonds | LINCS | 4 |

| Holo | | Step | Ensemble | Duration [$ns$] | Restraints [$kJ/mol$] Sidechain | Backbone | Lipid Z | Lipid dihed | Helix | CV | Continue | Repl |
|---|---|---|---|---|---|---|---|---|---|---|---|---|
| Initial AA-MD | Equil. | 0 | NVT | 0.125 (1fs) | 1000 | 2000 | 400 | 400 | – | – | No | 1 |
| | | 1 | NPT | 0.125 (1fs) | 500 | 1000 | 400 | 200 | – | – | Yes | 1 |
| | | 2 | NPT | 0.5 (2fs) | 200 | 500 | 200 | 200 | – | – | Yes | 1 |
| | | 3 | NPT | 1.0 (2fs) | 0 | 100 | 40 | 0 | – | – | Yes | 1 |
| | ABMD | 0 | NPT | COMITTOR (2fs) | 0 | 500$^a$ | 0 | 0 | 5000.0 | 2000.0$^b$ | Yes | 1 |
| | | 1 | NPT | 15.0 (2fs) | 0 | 20 | 200$^c$ | 0 | 0.0 | 0.0 | Yes | 1 |
| | Prod. | 0 | NPT | 60.0 (2fs) | 0 | 0 | 0 | 0 | – | – | No | 7 |
| | | 1 | NPT | 2000.0 (2fs) | 0 | 0 | 0 | 0 | – | – | Yes | 7 |
| CG-MD$^d$ | Equil. | 0 | NPT | 0.005 (1fs) | – | – | – | – | – | – | No | 9 |
| | | 1 | NPT | 1.250 (5fs) | 1000 | 1000 | – | – | – | – | Yes | 9 |
| | | 2 | NPT | 0.250 (10fs) | 1000 | 1000 | – | – | – | – | Yes | 9 |
| | | 3 | NPT | 0.500 (20fs) | – | – | – | – | – | – | Yes | 9 |
| | | 4 | NPT | 150.0 (20fs) | – | – | – | – | – | – | Yes | 9 |
| | Prod. | 0 | NPT | 2000.0 (2fs) | 0 | 0 | 0 | 0 | – | – | Yes | 9 |
| Re-seeded AA-MD$^e$ | Equil. | 0 | NVT | 0.125 (1fs) | 1000 | 2000 | 400 | – | 100 | – | No | 12 |
| | | 1 | NPT | 0.125 (1fs) | – | 2000 | 400 | – | 100 | – | Yes | 12 |
| | | 2 | NPT | 0.500 (2fs) | – | 1000 | 400 | – | 100 | – | Yes | 12 |
| | | 3 | NPT | 1.000 (2fs) | – | 100 | 40 | – | 100 | – | Yes | 12 |
| | | 4 | NPT | 20.00 (2fs) | – | – | – | – | 100 | – | Yes | 12 |
| | Prod. | 0 | NPT | 2000.0 (2fs) | 0 | 0 | 0 | 0 | – | – | Yes | 12 |

| Apo | Step | Ensemble | Duration [$ns$] | Restraints [$kJ/mol$] Sidechain | Backbone | Lipid Z | Lipid dihed | Helix | CV | Continue | Repl |
|---|---|---|---|---|---|---|---|---|---|---|---|
